# Fast and efficient generation of knock-in human organoids using homology-independent CRISPR/Cas9 precision genome editing

**DOI:** 10.1101/2020.01.15.907766

**Authors:** Benedetta Artegiani, Delilah Hendriks, Joep Beumer, Rutger Kok, Xuan Zheng, Indi Joore, Susana Chuva de Sousa Lopes, Jeroen van Zon, Sander Tans, Hans Clevers

## Abstract

CRISPR/Cas9 technology has revolutionized genome editing and is applicable to the organoid field. However, precise integration of exogenous DNA sequences in human organoids awaits robust knock-in approaches. Here, we describe <u>CRISPR</u>/Cas9-mediated <u>H</u>omology-independent <u>O</u>rganoid <u>T</u>ransgenesis (CRISPR-HOT), which allows efficient generation of knock-in human organoids representing different tissues. CRISPR-HOT avoids extensive cloning and outperforms homology directed repair (HDR) in achieving precise integration of exogenous DNA sequences at desired loci, without the necessity to inactivate TP53 in untransformed cells, previously used to increase HDR-mediated knock-in. CRISPR-HOT was employed to fluorescently tag and visualize subcellular structural molecules and to generate reporter lines for rare intestinal cell types. A double reporter labelling the mitotic spindle by tagged tubulin and the cell membrane by tagged E-cadherin uncovered modes of human hepatocyte division. Combining tubulin tagging with TP53 knock-out revealed TP53 involvement in controlling hepatocyte ploidy and mitotic spindle fidelity. CRISPR-HOT simplifies genome editing in human organoids.

---

Organoids can be generated by guided differentiation of induced pluripotent stem (iPS) cells and embryonic stem (ES) cells, or from cells isolated from adult tissues^1^. Adult stem cell (ASC)-derived organoids are self-organizing structures that recapitulate aspects of cellular composition, 3D architecture and functionality of the different epithelial tissues from which they originate, while maintaining genomic stability^2,3^. The possibility to derive organoids from genetically modified mouse lines, especially knock-in models, has allowed the generation of engineered mouse organoids, which have been used as versatile *in vitro* tools to answer various biological questions^4-10^.

Instead, the generation of engineered human ASC-derived organoids requires efficient strategies for *in vitro* genome editing to be applied after the lines have been established. CRISPR/Cas9 technology has considerably simplified genetic engineering. These approaches were so far largely limited to the NHEJ-mediated introduction of indels in the endogenous loci of organoids, leading to gene mutations^11,12^. By harnessing the homology directed repair (HDR) pathway, a single base substitution was introduced to correct the CFTR locus in cystic fibrosis intestinal organoids^13^, and a few human ASC-organoid knock-in reporter lines have been generated, but only in colon (cancer) organoids^14-16^. Knock-in by HDR takes advantage of a mechanism used by cells to repair double strand breaks (DSBs). Such breaks can be introduced at specific sites by CRISPR/Cas9. HDR is the most commonly used approach for targeted insertion, but this process is inefficient and requires cells to be in S phase^17,18^. Additionally, it requires cloning of the donor plasmid due to the necessity of the presence of homology arms specific to each gene (**Fig. 1a**). Recent studies have shown that CRISPR-induced DSBs activate the TP53 damage response and induce a transient cell cycle arrest in untransformed cells^19^. Permanent or transient inactivation of TP53 increases HDR-mediated knock-in in pluripotent and hematopoietic stem cells^20,21^. Thus, given the demand of novel ways to improve HDR efficiency, inhibition of TP53 was suggested as a potential solution to overcome the low HDR efficiency in untransformed cells^21^.

**Figure 1:**
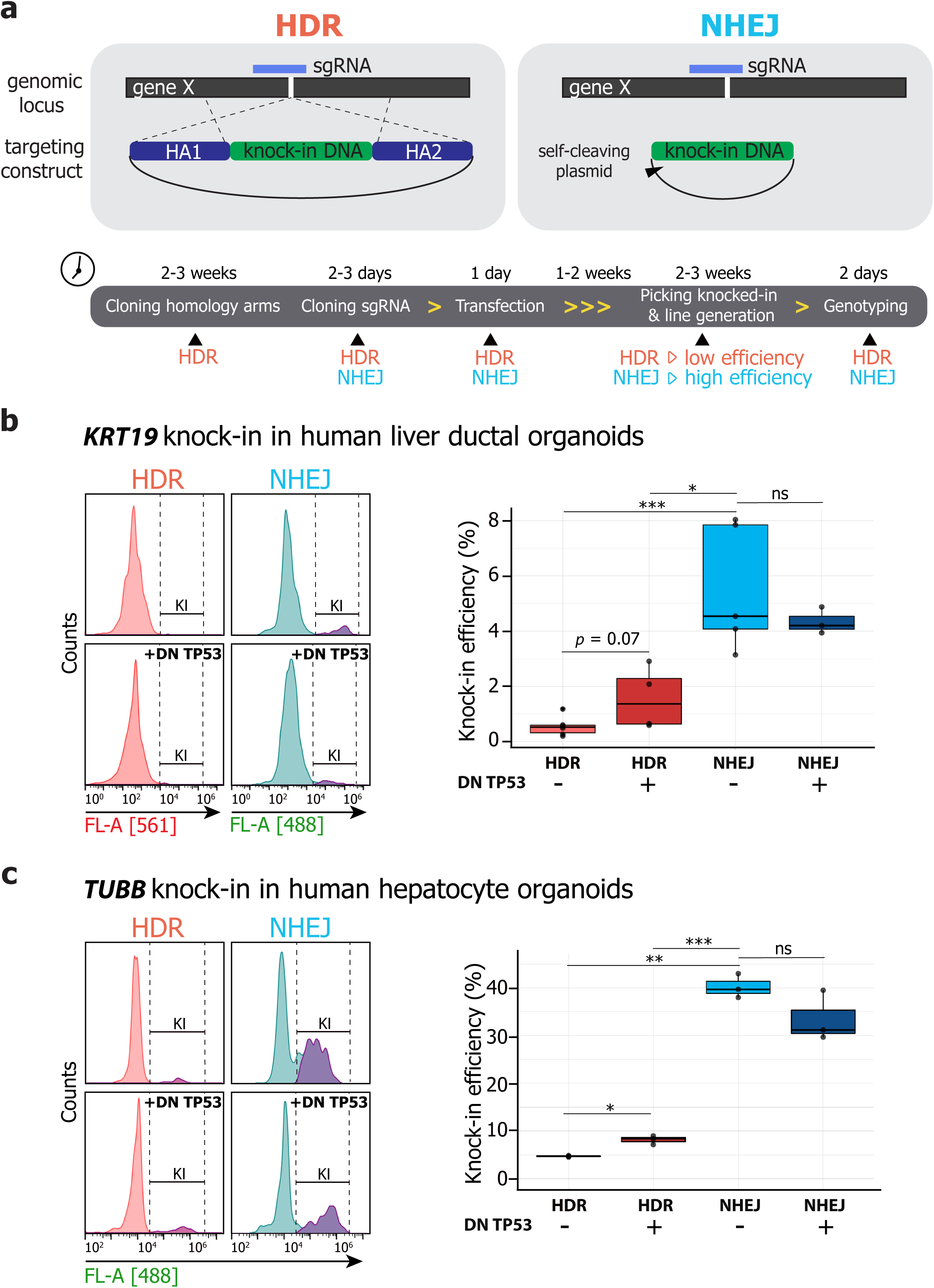
CRISPR-HOT strategy and efficiency. **(a)** Principle and step-wise description of using homology directed repair (HDR) vs. non-homologous end joining (NHEJ) to obtain knock-in of exogenous DNA into the genomic locus of interest and the associated timeline. Number of yellow arrowheads between steps are scaled to time. Note that for NHEJ, a targeting plasmid is used that is cleavable by a sgRNA at the 5’ of the knock-in tag. **(b)** Flow-cytometry histograms showing the knock-in (KI) efficiency at the *KRT19* locus in transfected human liver ductal cells using HDR (in red) vs NHEJ (in blue) with or without transient TP53 inhibition (by overexpression of a dominant negative form of TP53 (DN TP53)). Quantification of the percentage of knocked-in cells over the transfected cells in the different conditions is shown on the right. Data are represented as the mean ± s.d. of knock-in efficiencies in organoid lines from at least 3 different donors assessed in independent experiments. **(c)** Comparison of the knock-in efficiency at the *TUBB* locus in transfected human hepatocytes using HDR or NHEJ with or without transient TP53 inhibition as determined by FACS analysis as in **b**. Data are represented as the mean ± s.d. of knock-in efficiencies of 3 independent experiments.

Non-homologous end joining (NHEJ), another key DNA repair system, is active in all cell cycle phases^18^, and by ligating DNA ends, does not require regions of homology (**Fig. 1a**). Since it is generally believed to be error prone, NHEJ is not widely used for precision transgene insertion. Yet, it has been suggested that NHEJ can be fundamentally accurate and can re-ligate DNA ends without mistakes^22,23^. Indeed, a handful of studies have exploited NHEJ to ensure the targeted insertion of exogenous DNA in zebrafish^24^, mouse^25^, immortalized human cell lines^26,27^, and ES cells^28^. Here, we leverage NHEJ-mediated knock-in to the human organoid field, an approach termed CRISPR-HOT, as a versatile, efficient and robust homology-independent method to obtain knock-in human wild-type organoids from different organs.

## Results

### CRISPR-HOT is a novel strategy to efficiently generate knock-in human organoids

We first focused on two different -and hard to transfect-human organoid systems from the two main epithelial cell types of the liver (human liver ductal^29^ and hepatocyte^30^ organoids). We i) tested the efficiency of the HDR conventional approach, ii) assessed whether NHEJ can be used for precise genome editing and knock-in of exogenous DNA at specific genomic loci, and iii) performed a side-by-side comparison of the two strategies. For HDR, we employed donor plasmids containing a fluorescent tag (tdTomato or mNEON) flanked by 0.5 kb homology arms to C-terminal target *KRT19* or *TUBB* in human liver ductal organoids and human hepatocyte organoids, respectively. For NHEJ, we utilized a universal targeting plasmid containing a fluorescent protein, in this case mNEON, for C-terminal gene tagging (see **Fig. 1b** and **Supplementary Fig. 1** for an overview and schematic representation of all targeting plasmids used in this study). This plasmid is cleavable due to the upstream incorporation of a non-human sequence which can be recognized by a sgRNA, thus mediating linearization of the plasmid by CRISPR/Cas9^27^. To promote in-frame knock-in, this non-human sequence can be targeted by three different sgRNAs (frame selectors). Each of these frame selectors mediates the cut in one of the three possible frames. This strategy has been previously employed for immortalized human cell lines^26,27^.

Liver ductal organoids are composed of a simple epithelium, with cells expressing early cholangiocyte markers such as KRT19^29^. To test HDR-mediated knock-in at the *KRT19* locus, we transfected human liver ductal organoids with the HDR targeting plasmid, a plasmid expressing Cas9 and a fluorescent marker as transfection readout, and a plasmid expressing a sgRNA inducing a DSB directly upstream of the *KRT19* stop codon, using our recently described method^31^. To test NHEJ, we employed the same transfection approach using instead the universal NHEJ self-cleaving targeting plasmid, the same KRT19 sgRNA plasmid and the appropriate frame selector, which also expresses Cas9 and a fluorescent marker (mCherry) as transfection readout. Experiments were repeated using organoid cultures from five different donors. Five days after transfection, we observed some knocked-in cells in the HDR-condition (0.55% ± 0.35%) as measured by FACS analysis of knocked-in cells within the transfected population (transfection efficiency ranging from 0.2-0.7%). Instead, significantly more knocked-in cells appeared when testing NHEJ-mediated gene insertion (5.53% ± 2.26%) (**Fig. 1b, Supplementary Fig. 2a-c**).

**Figure 2:**
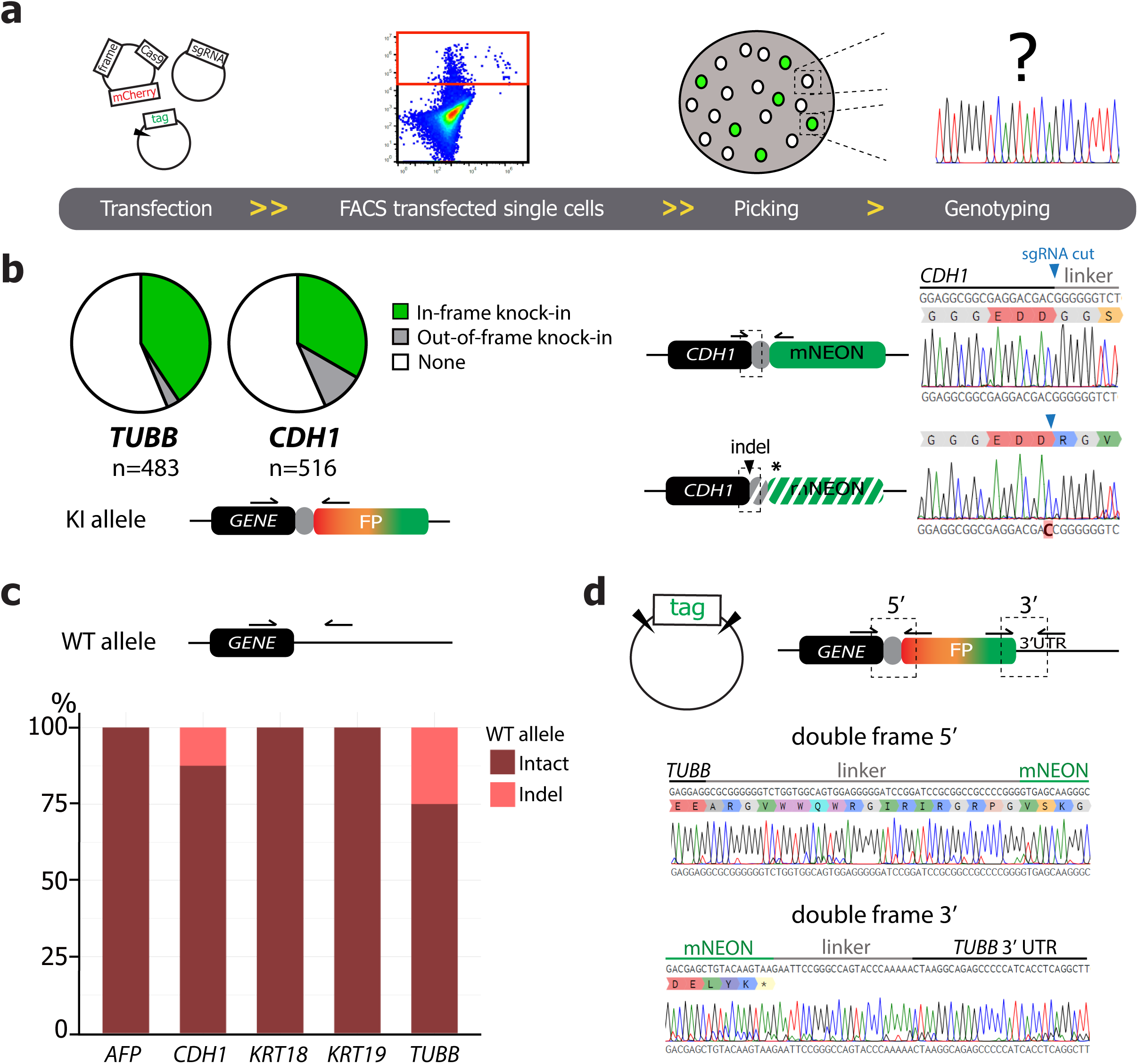
CRISPR-HOT allows precise gene knock-in. **(a)** Schematic overview of the experimental set-up to assess the preciseness of NHEJ-mediated knock-in events. Human hepatocyte organoids were transfected with the NHEJ plasmids, single transfected cells were FAC-sorted, and outgrowing clonal organoids (both mNEON^+^ and mNEON^-^) were sequenced. **(b)** Sequencing of the PCR-amplicons using primers spanning the region of knock-in shows that CRISPR-HOT mediates predominantly precise in-frame knock-in at two different loci (*TUBB* and *CDH1*) in human hepatocyte organoids (left). Representative sequencing examples of in-frame and out-of-frame knock-in events at the *CDH1* locus (right). **(c)** Analysis of the untargeted allele from clonal and bulk organoid lines knocked-in at 5 different loci. Bar plot showing the percentage of carrying indels or being intact resulting from 9 independent targeting experiments. **(d)** A double frame selector allows cleaving of the targeting plasmid at both the 5’ and 3’ of the tagging cassette. This plasmid was tested at the *TUBB* locus. Sequencing of both 5’ and 3’ ends demonstrate precise knock-in of this excised tag at both ends. Sequencing examples representative of multiple clonal organoids are shown.

We performed similar experiments (in triplicate) in human hepatocyte organoids targeting another locus, *TUBB*. For this organoid system, we adapted a different transfection method relying on cuvette electroporation. In this experimental set-up, we again observed a considerably higher efficiency of NHEJ (40.23% ± 2.54%) as compared to HDR-mediated knock-in (4.70% ± 0.15%) within the transfected population (transfection efficiency ranging from 9-14%) (**Fig. 1c, Supplementary Fig. 2a, b**). Outgrowing knocked-in organoids from both conditions displayed similar localization of the fluorescent signal, characteristic of TUBB expression patterns (**Supplementary Fig. 2e**). Having validated that NHEJ can mediate knock-in in human organoids we termed this approach CRISPR-HOT: <u>CRISPR</u>/Cas9-mediated <u>H</u>omology-independent <u>O</u>rganoid <u>T</u>ransgenesis.

Recently, different studies have shown that transient TP53 inhibition can improve HDR efficiency in human pluripotent stem cells^20^ and hematopoietic stem cells^21^. Therefore, we checked how the efficiency of this published strategy compared to the efficiency of CRISPR-HOT, by performing a direct comparison of HDR and CRISPR-HOT in presence of transient TP53 inhibition by co-transfection of a dominant negative form of TP53 (DN TP53)^20,32^. Although the presence of the DN TP53 increased HDR efficiency both in human liver ductal (1.56% ± 1.13%) and hepatocyte (8.18% ± 0.93%) organoids (**Fig. 1b-c**), it still was only 2-fold higher than HDR in the absence of DN TP53 and much lower than the efficiencies reached with NHEJ (10-fold higher compared to HDR) (**Supplementary Fig. 2d**). DN TP53 did not affect NHEJ-mediated knock-in efficiency (**Fig. 1b-c**). Of note, while absolute knock-in efficiencies between the two organoid systems was variable, relative fold-changes between HDR, NHEJ, and the presence or absence of DN TP53 were remarkably similar (**Supplementary Fig. 2d**). Thus, these results demonstrated that our homology-independent genome editing approach CRISPR-HOT can mediate DNA insertion in human organoids at higher efficiency than previously used methods.

### CRISPR-HOT mediates precise gene insertion

Having assessed the capacity of CRISPR-HOT to mediate gene tagging in human organoids, we next evaluated how frequently the DNA integration is precise. The frame-selector strategy previously explained should increase the proportion of desired in-frame knock-in events. To test this, we transfected human hepatocyte organoids with the plasmids required to target two different loci, *TUBB* and *CDH1*, in independent experiments. After FAC-sorting of transfected mCherry^+^ cells, we quantified the proportion of mNEON^+^ and mNEON^-^ organoids based on fluorescent imaging of a 24-well for each condition. We then picked individual clonal organoids outgrowing from these single cells and sequenced both knocked-in mNEON^+^ organoids as well as mNEON^-^ organoids (**Fig. 2a**). In case of targeting *CDH1*, 33% of the organoids had precise in-frame knock-in; for *TUBB* this proportion was 40%, which was at levels very comparable to our previous observation of knock-in efficiency at this locus (**Fig. 2b**, and compare with **Fig. 1c**). This also indicates that the vast majority of knocked-in cells are able to grow out as organoids. Analysis of the mNEON^-^ organoids revealed a small percentage of imprecise out-of-frame knock-in events due to the introduction of small indels (*CDH1*: 10%; *TUBB*: 3% of the total population), while the rest did not have insertions (both *CDH1* and *TUBB* 56% of the total population) (**Fig. 2b**). Two representative sequencing examples of in-frame and out-of-frame knock-in at the *CDH1* locus are given in **Fig. 2b**.

Next, since the introduction of a Cas9-mediated DSB could generate indels in the allele that was untargeted by the knock-in event, we PCR-amplified and sequenced a DNA region around the cut site. By analysing five different loci targeted across two different human organoid systems (liver ductal and hepatocyte) in 9 independent experiments using both bulk and clonal organoid cultures, indels were found at only two loci (*CDH1*: 12.5%; *TUBB*: 25%) (**Fig. 2c**). Thus, it is possible to select for the desired genotype on both alleles based on sequencing, and the likelihood to obtain the desired genotype is high. While in the case of C-terminal gene tagging, potential indels in the untargeted allele should not interfere with gene function, this is important to consider for N-terminal gene tagging. We constructed a double frame selector strategy, where the tag of interest is flanked by two frame selector cassettes on both 5’ and 3’ sides, which can be targeted by the same sgRNA so that the tag fragment can be excised and incorporated into the genomic region of interest in-frame as well as maintaining the reading frame of the remaining 3’ of the targeted gene (**Fig. 2d**). As a proof of concept, we validated this approach by targeting *TUBB* with this double frame plasmid in human hepatocyte organoids. We observed mNEON^+^ organoids and sequencing of multiple independent clonal organoids revealed precise in-frame insertion at both the 5’ and 3’ sides of the inserted mNEON tag (**Fig. 2d**). This double frame selector strategy is an important asset that could be obviously also potentially applied to N-terminally tag genes. Altogether, CRISPR-HOT can mediate precise in-frame insertion with high efficiency while maintaining the second allele intact.

### Generation of human liver ductal reporter organoid lines by CRISPR-HOT

We next evaluated whether we could derive clonal organoid lines from the KRT19-tagged human liver ductal cells. Two days after electroporation, transfected organoids were picked based on transient mCherry expression from the frame selector plasmid. These were subsequently dissociated into single cells to grow clonal lines^31^ (**Fig. 3a**). Single knocked-in mNEON^+^ organoids were picked and grown out as KRT19::mNEON clonal lines. Correct insertion at the *KRT19* locus was validated by sequencing (**Fig. 3b, c**). KRT19 is expressed in all cells in human liver ductal organoids^29^. Accordingly, mNEON expression was observed in every cell in KRT19::mNEON organoids. Staining for ZO-1 and actin confirmed that these knock-in organoids showed robust epithelial polarization typical for human liver ductal organoids (**Fig. 3c**). Cytoplasmic cellular distribution was also in agreement with the expected KRT19 localization by antibody co-staining (**Fig. 3d**).

**Figure 3:**
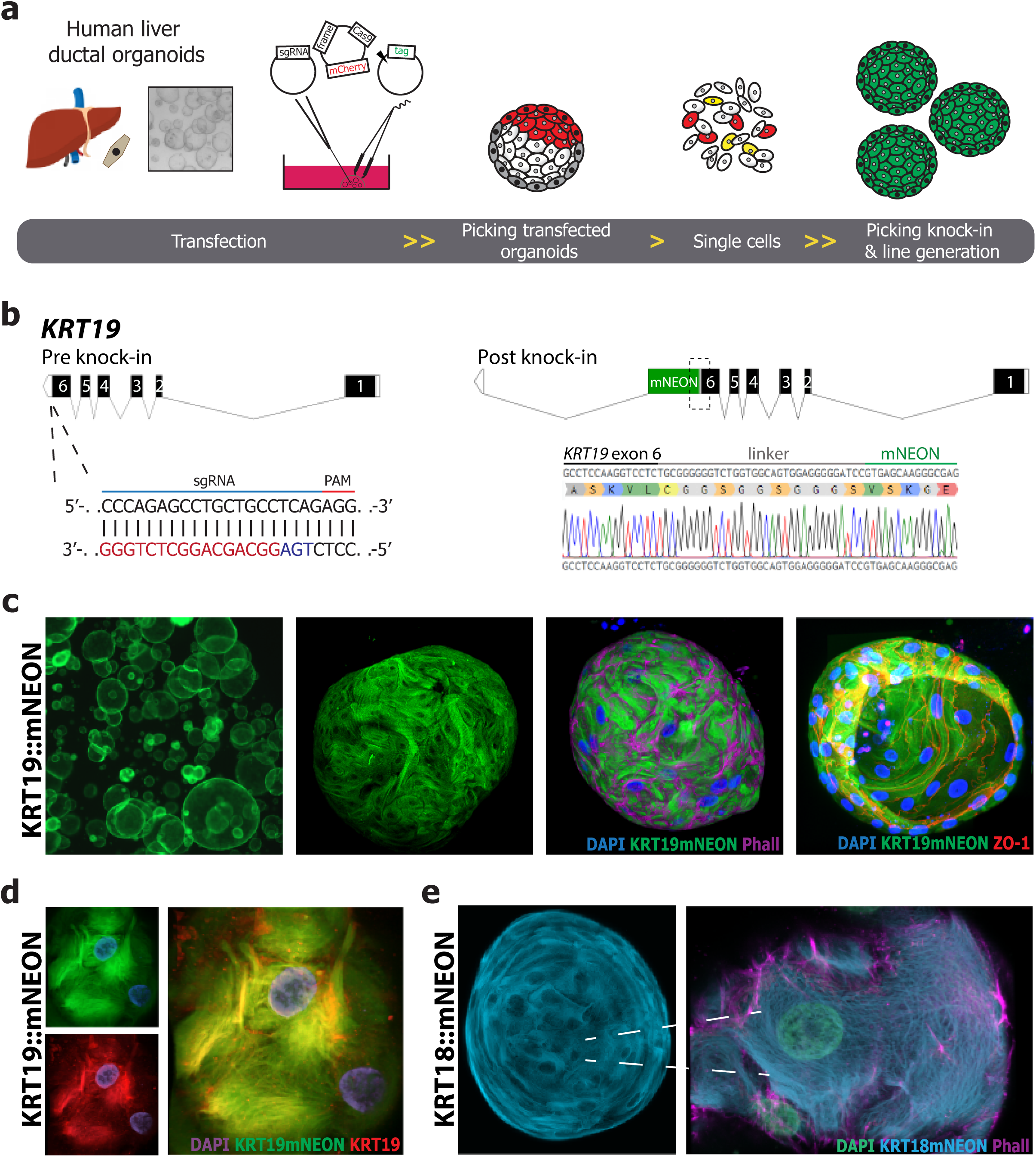
Generation of human liver ductal reporter organoid lines using CRISPR-HOT. **(a)** Transfection strategy for human liver ductal organoids. DNA is microinjected into the lumen of the organoids grown in a BME^R^ droplet using a glass capillary, where after the BME^R^ droplet is electroporated using a tweezer electrode. Organoids with transfected cells are picked based on transient mCherry expression from the frame selector plasmid and made into single cells. In-frame knocked-in mNEON^+^ organoids growing from these single cells are subsequently picked to establish clonal reporter lines. **(b)** Strategy and sgRNA used to tag the human *KRT19* locus. Schematic overview of the *KRT19* locus pre- and post knock-in. A representative sequencing result confirms in-frame knock-in of mNEON at the *KRT19* locus. **(c)** Representative fluorescent image of a clonal KRT19::mNEON culture at low and high-magnification (left). Whole-mount of a KRT19::mNEON organoid stained with Phalloidin and ZO-1 (right). **(d)** Co-staining of KRT19::mNEON with a KRT19 antibody demonstrates overlapping signals. **(e)** Representative fluorescent image of a clonal KRT18::mNEON culture (left). High magnification imaging of a KRT18::mNEON organoid. The endogenous KRT18mNEON fusion allows visualization of the intermediate filament structures, while the actin cytoskeleton is visualized by Phalloidin (right). See **Supplementary Fig. 3a** and **Fig. 8a** for a schematic overview of the *KRT18* locus pre- and post knock-in and representative sequencing results of the KRT18::mNEON line.

Another keratin, KRT18, was targeted as a second locus to validate the robustness of our approach. Again, precise in-frame integration was obtained when targeting this locus (**Supplementary Fig. 3a**). Expression of mNEON in the KRT18::mNEON line was confined to the cytoplasm and high-resolution imaging allowed labelling of intermediate filaments (**Fig. 3e**). Altogether, our data showed that hard-to-transfect human liver organoids^33^ can be genome-edited to generate clonal knock-in reporter lines using CRISPR-HOT.

### CRISPR-HOT enables labelling of rare intestinal cell types

We next tested whether CRISPR-HOT also works in human organoids from another organ. Human intestinal organoids can be induced to contain several differentiated cell types, mirroring the cell types present in the intestine *in vivo*^34,35^. In contrast to mouse intestinal organoids, differentiation of human intestinal organoids is less efficient and requires a change in culture conditions^8,35^. Rare intestinal cell types, such as hormone-producing enteroendocrine cells, are therefore relatively sparse when compared to mouse intestinal organoids. Yet, generation of reporter lines to identify these human cell types can aid studies to understand their functions and regulation.

We evaluated whether intestinal organoids could also be genome-edited using CRISPR-HOT. To this end, we adapted a different transfection method based on cuvette electroporation of small clumps of cells^36^ and FAC-sorting after 5-7 days (**Fig. 4a**). We decided to target a broadly expressed gene (*CDH1*) as well as two marker genes expressed in specific and rare cell types (*CHGA* for enteroendocrine cells and *MUC2* for goblet cells). For the constitutively expressed *CDH1*, we could directly pick clonal knocked-in organoids that grew out after FAC-sorting (**Fig. 4a**). Instead, for the non-constitutively expressed, differentiated marker genes (*CHGA* and *MUC2*), we used a short differentiation pulse (specific to the desired cell population)^8,37^ which could be reversed by switching back culture conditions to expansion medium. This approach allows visualization of positive cells within one knocked-in organoid derived from one single FAC-sorted cell, which can then be subsequently picked to establish clonal knock-in organoid lines (**Fig. 4a**). Accordingly, when we targeted the *CDH1* locus, we could already identify knocked-in mNEON^+^ cells within the transfected cells (transfection efficiency ranging from 1.6-2.6%) during FAC-sorting (**Fig. 4b**). In contrast, as expected, we did not observe any mNEON^+^ cells when targeting the *CHGA* or *MUC2* locus (see the *CHGA* example in **Fig. 4b**) as fluorescent signals can only be visualized upon induction of gene expression related to the differentiation into the respective cell type.

**Figure 4:**
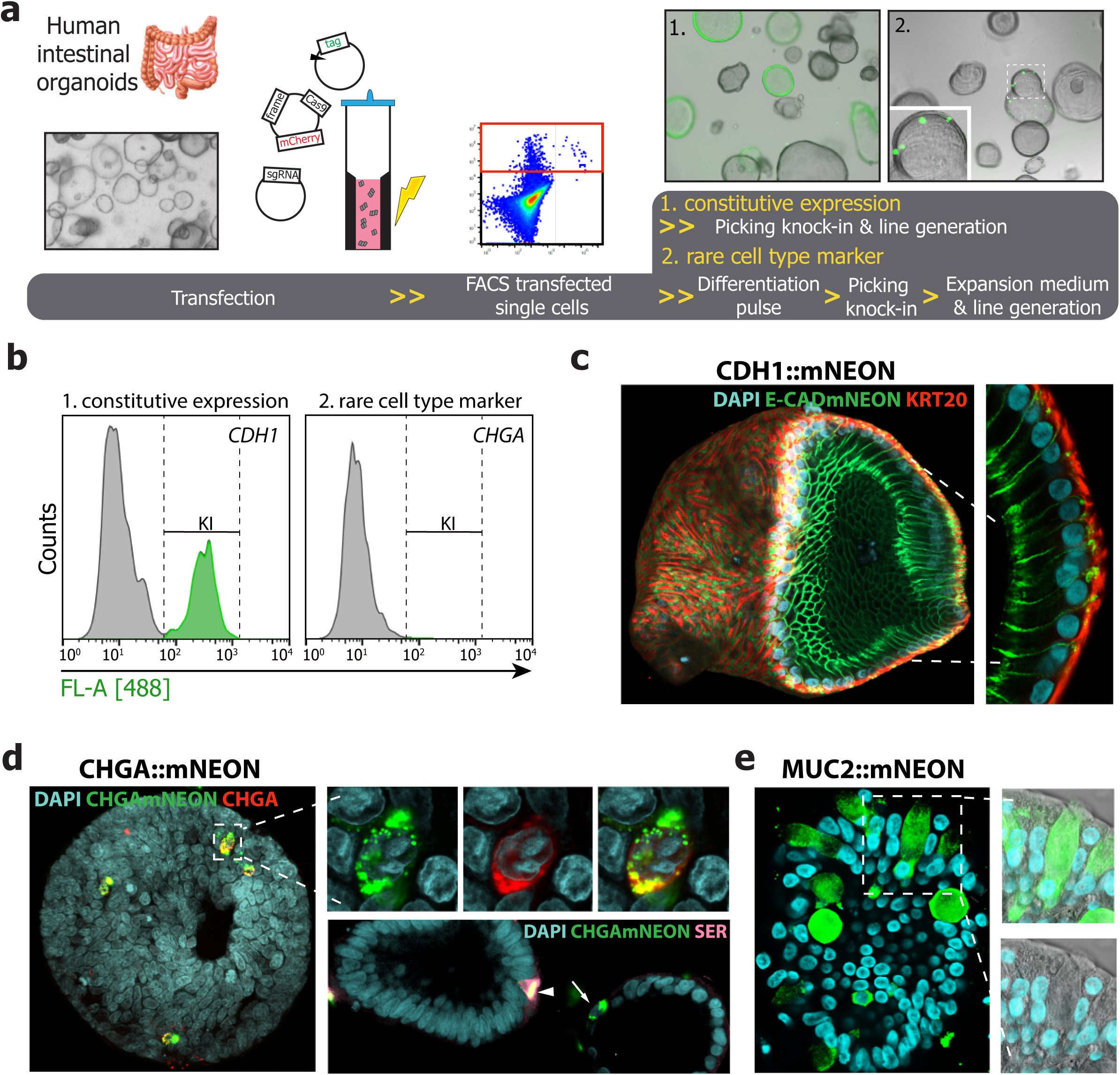
Labelling of different human intestinal cell types in organoids using CRISPR-HOT. **(a)** Transfection strategy for human intestinal organoids. Organoids are dissociated into small clumps of cells, mixed with DNA, and electroporated in a cuvette. Single transfected cells are FAC-sorted and grown into organoids. When tagging constitutively expressed genes, single clonal tagged organoids are picked and used to establish clonal lines (1). For tagging of non-constitutively expressed differentiation markers a reversible differentiation pulse with appropriate differentiation medium is used to visualize knocked-in organoids by the appearance of a few positive cells, which are subsequently picked and expanded in expansion medium to grow clonal lines (2). **(b)** Representative flow-cytometry histograms showing sorting of transfected cells in independent experiments of targeting *CDH1* and *CHGA*. Note that while for *CDH1* knocked-in cells are already visible during FAC-sorting, for *CHGA* no positive knocked-in cells are observed at this stage, since a differentiation pulse is required. **(c)** Representative whole-mount image of a CDH1::mNEON organoid differentiated towards enterocytes and co-stained with KRT20. **(d)** Whole mount staining of a CHGA::mNEON intestinal organoid for enteroendocrine cells stained for CHGA and Serotonin (SER). Arrowhead indicates a CHGA^+^SER^+^ cell while the arrow points at a CHGA^+^SER^-^ cell (middle). Representative transmitted and fluorescent image of a MUC2::mNEON intestinal organoid reporter for goblet cells (right). See **Supplementary Fig. 3b-c** and **Fig. 8a** for a schematic overview of the *CDH1* and *CHGA* loci pre- and post knock-in and representative sequencing results of these lines.

The mNEON^+^ *CDH1*-tagged human intestinal organoids (28.6% ± 0.1% knock-in efficiency) could be picked and readily grown into CDH1::mNEON clonal lines. E-CADmNEON showed typical membrane expression of all KRT20^+^ enterocytes (**Fig. 4c**). For *CHGA*, we picked single knock-in organoids (20.2% ± 5.0% knock-in efficiency) based on the short differentiation pulse using previously described enteroendocrine cell enrichment medium^8^, and established CHGA::mNEON clonal organoid lines. Upon expansion and subsequent differentiation, these organoids contained rare enteroendocrine (CHGAmNEON^+^) cells with fluorescently marked vesicles, *i.e.* secretory granules (**Fig. 4d**). These cells stained for CHGA protein, while a proportion also produced the neurotransmitter serotonin (**Fig. 4d**). For *MUC2*, we used a similar approach, but using a differentiation pulse based on addition of the Notch inhibitor DAPT^37^. Clonal organoid lines could be established and -upon differentiation-contained cells expressing MUC2mNEON showing the typical morphology of goblet cells (**Fig. 4e**). CRISPR-HOT appeared to be robust since we could efficiently derive clonal lines in multiple independent experiments for each gene targeted. In all cases, the derived lines showed precise in-frame knock-in, as confirmed by sequencing (see for example **Supplementary Fig. 3b, c**).

### Generation of clonal human hepatocyte reporter organoid lines using CRISPR-HOT

We already showed that CRISPR-HOT can be used to obtain knocked-in human hepatocytes (**Fig. 1c**). We next evaluated whether these could be picked to establish clonal lines. Organoids were first dissociated into single cells and then electroporated in a cuvette (**Fig. 5a**). We picked the knocked-in organoids that grew after transfection based on fluorescence and established clonal lines. First, we successfully targeted a gene exclusively expressed in hepatocyte organoids by virtually all cells, *AFP*^30^ and generated clonal AFP::mNEON lines (**Fig. 5b, Supplementary Fig. 3d**). AFP::mNEON organoids showed typical hepatocyte morphological features, such as the formation of bile canalicular networks, as visualized by MRP2 staining (**Fig. 5b**). AFP in these organoids had the same pattern of expression visualized by antibody staining, yet the AFPmNEON signal has higher signal resolution as compared to the antibody that is uncapable of fully penetrating the organoids (**Fig. 5b**).

**Figure 5:**
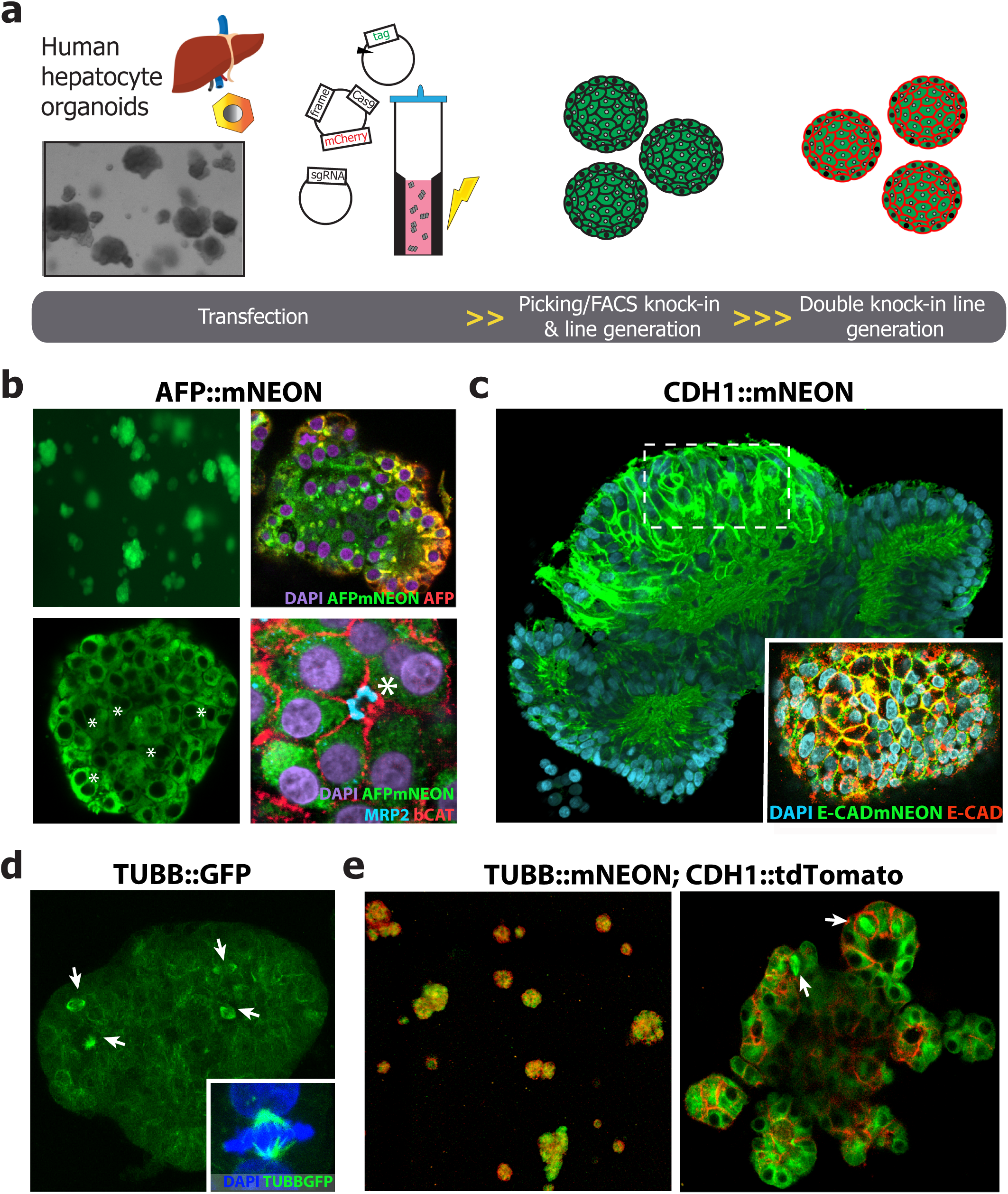
Generation of human hepatocyte reporter organoid lines using CRISPR-HOT. **(a)** Transfection strategy for human hepatocyte organoids. Organoids are dissociated into small clumps of cells, mixed with DNA, and electroporated in a cuvette. Single transfected cells grow into organoids, which can be picked to establish clonal lines. An additional locus can be eventually targeted using the same procedures to establish double knock-in reporters. **(b)** Representative image of a clonal AFP::mNEON line. Co-staining of AFP::mNEON with an AFP antibody demonstrates overlapping signal but limited penetration of the antibody. Note the typical hepatocyte polarity visualized by staining for the bile canalicular marker MRP2, while the cell membrane is marked by β-catenin. Asterisks indicate the presence of polyploid (binucleated) hepatocytes. **(c)** Representative image of a CDH1::mNEON hepatocyte organoid. In the inset, staining for E-cadherin demonstrates perfect overlap with E-CADmNEON. **(d)** Representative image of a TUBB::GFP hepatocyte organoid. Inset shows a dividing cell with the mitotic spindle visualized by TUBB::GFP and chromatin by DAPI staining **(e)** Representative image of a clonal TUBB::mNEON; CDH1::tdTomato double knock-in hepatocyte line (left) and high magnification imaging of a respective organoid (right). In **d-e** arrows indicate mitotic events. See **Supplementary Fig. 3d-f** and **Fig. 8a** for a schematic overview of the *AFP, TUBB*, and *CDH1* loci pre- and post knock-in and representative sequencing results of these lines.

Similar to the intestinal organoids, we were able to derive clonal CDH1::mNEON hepatocyte lines (**Fig. 5c**). The presence of the endogenously tagged *CDH1* allowed to trace cell movement and dynamics. For instance, time-lapse movie analyses revealed that hepatocytes tend to organize around a central lumen and rotate around it, while forming “rosette structures” in the organoids (**Supplementary Fig. 4**). These observations are based on analyses performed in *CDH1*-tagged human hepatocyte organoid lines derived from two different donors.

We also established clonal *TUBB*-tagged hepatocyte organoids as a tool to be able to trace real-time mitotic spindle dynamics of proliferating hepatocytes (**Fig. 5d, Supplementary Fig. 3e**). Being able to generate lines with simultaneous tagging of more than one locus, could be a major advantage for tracking detailed and short-lived cellular changes. Therefore, since concomitant visualization of the cell membrane and mitotic spindle would allow more refined analysis of division events, we decided to sequentially tag *CDH1* with tdTomato in a TUBB::mNEON line (**Fig. 5a**). Indeed, we could derive double knocked-in TUBB::mNEON;CDH1::tdTomato clonal lines (**Fig. 5e, Supplementary Fig. 3f**).

### Mitotic spindle analyses reveal an orchestrated mode of hepatocyte division

Hepatocytes in the liver are organized in a peculiar fashion reflecting features of stratified as well as monolayered (simple) epithelia^38^. Mitotic spindle positioning and the establishment of proper polarity and tissue organization are known to be linked, although how this exactly works in the liver is still debated^39,40^. A recent study underscored that tissue architecture ensures correct mitotic spindle behaviour. Indeed, 2D cultures of dissociated epithelial cells display mitotic spindle defects opposite to what is observed in cells cultured in a 3D configuration^41^. Human hepatocyte organoids retain cellular polarization, with an apical domain defined by expression of MRP2 and ZO-1, facing the internal bile canaliculus (**Fig. 5b** and^30^). Our TUBB-tagged lines give us the possibility to perform analysis of non-transformed human hepatocytes within a 3D structure. We performed time-lapse analyses (**Supplementary Video 1**) and measured changes of spindle position as defined by the angle of rotation of spindle axis over time from metaphase to telophase (**Fig. 6a, Supplementary Fig. 5a**). We noticed that the spindle position is not static, but instead can vary considerably. In particular, we observed that spindles that have a similar position in prometaphase and cytokinesis (arbitrarily defined as a difference of less than a 45-degree angle between the two stages) hardly rotate (**Fig. 6b**, top). In contrast, spindles that change their axis between prometaphase and cytokinesis tend to rotate considerably over time (**Fig. 6b**, bottom), possibly trying to reach a specific orientation. In addition, we measured the spindle position over time relative to the apical domain which could be visualized by the presence of internal lumina. For this analysis, we defined the spindle axis and considered the angle that it forms with the apical side (**Fig. 6c-d**). While the orientation of the spindle appeared to be random at prometaphase and during the following mitotic stages, before cytokinesis it consistently ended up within an angle between 0 and 45-degrees relative to the apical side (**Fig. 6e**). Importantly, these conclusions were derived from multiple independent experiments using two donors displaying similar behaviour. Taken together, these observations suggest that at telophase the position of the spindle is not random and that hepatocytes dividing next to a lumen undergo symmetric division with respect to the apicobasal axis, as confirmed by time-lapse analysis of cell division in the CDH1::mNEON line (**Fig. 6f**).

**Figure 6:**
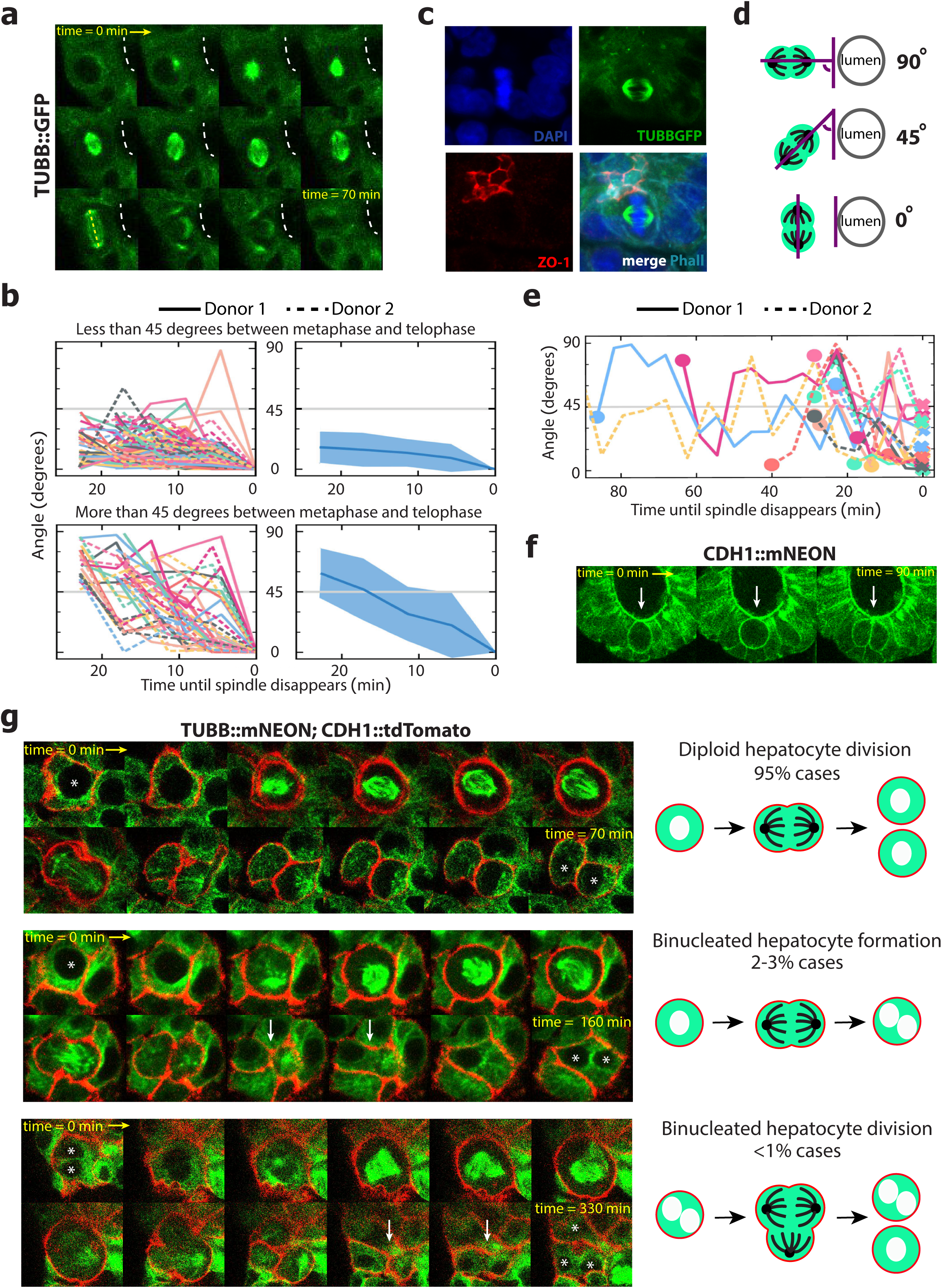
Mitotic spindle analyses in human hepatocyte reporter organoids reveal division dynamics. **(a)** Time-lapse images (5 min interval) of a TUBB::GFP organoid visualizes a hepatocyte dividing next to lumen (white dashed line). While the spindle initially rotates, it ends up parallel (yellow dashed line) to the apical luminal side, resulting in symmetric cell division. **(b)** Analysis of the spindle rotation in hepatocytes from metaphase to telophase. The upper panel shows mitotic spindles with a starting angle differing less than 45 degrees from its final angle; the lower panel shows spindles with an angle difference of more than 45 degrees. The spindle angle is defined by the spindle axis and is relative to the final angle (defined as 0 degrees). Each line represents 1 mitotic spindle, with mitosis being analysed from 2 different donors. **(c)** Staining for the apical marker ZO-1 in TUBB::GFP organoids confirms parallel alignment of the mitotic spindle with the apical side in telophase. **(d)** Schematic representation of how possible orientations of the mitotic spindle position at telophase relative to the lumen were defined. **(e)** Analysis of the spindle rotation of hepatocytes relative to the apical side as visualized by the presence of a lumen demonstrates that spindles tend to align with the apical side upon exiting telophase. Each line represents 1 mitotic spindle, with mitosis being analysed from 2 different donors. Circles on the left represent metaphase (initial orientation) and crosses on the right telophase (final orientation). **(f)** Time-lapse imaging (45 min interval) of CDH1::mNEON organoids reveals that hepatocyte division occurs at the apical side and results in symmetric cell division. **(g)** Characterization of mitotic events in the double TUBB::mNEON;CDH1::tdTomato hepatocyte reporter line. Representative snapshots of a time-lapse imaging experiment (5 min interval), showing diploid hepatocyte division (top), binucleated hepatocyte formation (middle), and binucleated hepatocyte division (bottom). Each asterisk indicates a nucleus. White arrows point at the E-CADtdTomato membrane forming transiently in the process of forming a binucleated cell.

The liver is characterized by the presence of polyploid hepatocytes, cells with more than two sets of chromosomes. These cells generally tend to increase in number with age and upon injury ^42^. Past studies have suggested that polyploid hepatocytes predominantly originate from incomplete cytokinesis^43,44^. Our double TUBB::mNEON;CDH1::tdTomato hepatocyte knock-in line gave us the possibility to examine polyploid events due to the simultaneous visualization of both the mitotic spindle and the cell membrane (**Supplementary video 2**). The vast majority of mitotic events (>95%) originated from diploid hepatocyte division (**Fig. 6g**). In a small percentage (2-3%), we observed the formation of a binucleated cell from a division of a mononucleated cell. Interestingly, while the daughter cells appeared segregated as marked by an initial formation of an ECADtdTomato^+^ cell membrane separating the two, this transient border disappeared resulting in a binucleated cell (**Fig. 6g**). We observed extremely rare (<1%) mitotic events encompassing division of binucleated cells through a three-polar spindle. Division resulted in the formation of a mononucleated and binucleated cell (**Fig. 6g**). Also in these cases, the formation of the binucleated cell was preceded by a transient ECADtdTomato^+^ border between two of the daughter cells (**Fig. 6g**). In addition, we did note the formation of monopolar spindles, but such cells did not complete division and eventually died, as also visualized by live imaging with a viability dye, and staining for cleaved caspase-3 (**Supplementary Fig. 5b-d**).

### Loss of TP53 induces aberrant mitotic spindle behaviour in human hepatocytes

TP53 is critical to maintain genomic stability and loss of TP53 is associated with increased polyploidy and aneuploidy^45,46^. Yet, its specific role in mitotic spindle behaviour is not fully understood. Here we combined CRISPR-HOT with CRISPR/Cas9-mediated gene knock-out to reveal the impact of loss of TP53 in our TUBB-tagged human hepatocyte lines. To this end, we transfected these cells with a plasmid that co-expresses Cas9 and a sgRNA targeting exon 4 of *TP53*^11^. TP53-mutated organoids were picked based on Nutlin-3a resistance, clonally expanded, and subsequent sequencing revealed the presence of frameshift indels (**Fig. 7a**). When performing time-lapse analyses on these clonal lines, we frequently observed hepatocyte divisions associated with the formation of non-canonical mitotic spindles (*i.e.* different from the canonical bipolar mitotic spindle) (**Fig. 7b-f, Supplementary Fig. 6**, and **Supplementary Video 3**). In particular, we detected an increase in multipolar spindle formation that appeared to result from the division of hepatocytes with increased ploidy (**Fig. 7c, Supplementary Fig. 6**). In contrast to wild-type hepatocyte organoids, in these *TP53*-deficient organoids we observed the presence of polyploid hepatocytes with increased nuclei (3-4) per cell, which retained the capacity to divide (**Fig. 7c**). Moreover, microtubules appeared frequently disorganized, for example leading to a transient collapse of the mitotic spindle, yet these cells would still continue to divide (**Fig. 7d, Supplementary Fig. 6**). We observed these aberrations in multiple clonal *TP53*^-/-^;TUBB-tagged hepatocyte lines from two different donors (3 clones per donor). Of note, division of hepatocytes with these non-canonical mitotic spindles was not associated with loss of the internal lumina, neither did the loss of TP53 change proliferation rates (**Supplementary Fig. 7a-c**). Altogether, these results imply that TP53 is of critical importance for mitotic spindle integrity and its loss is associated with the formation and division of polyploid hepatocytes.

**Figure 7:**
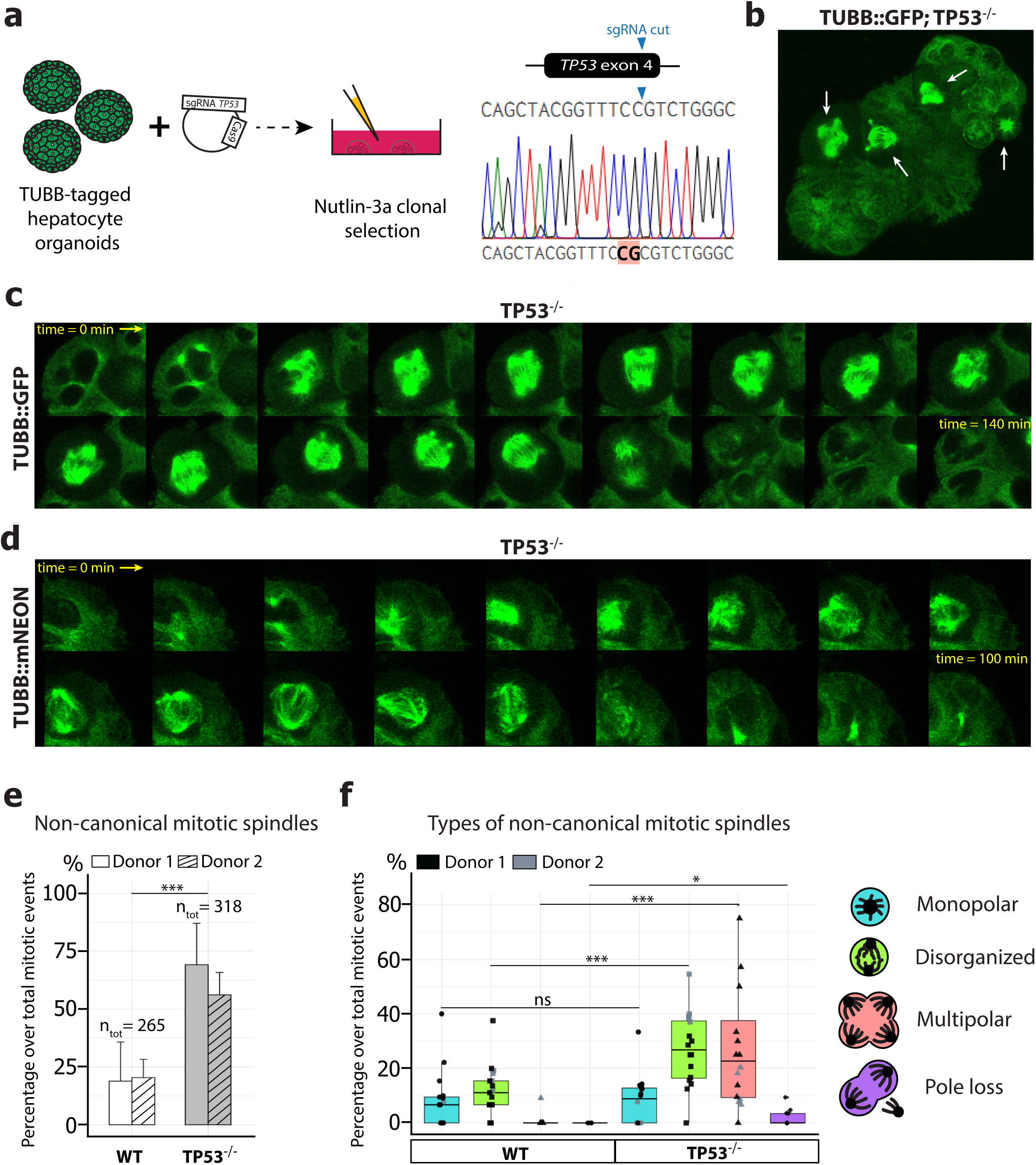
Loss of TP53 causes mitotic spindle aberrations in human hepatocytes. **(a)** Strategy of mutating *TP53* in TUBB-tagged hepatocyte organoids. Organoids are co-transfected with a plasmid carrying Cas9 and a sgRNA targeting TP53 and mutated organoids are selected based on Nutlin-3a resistance. A representative sequencing result from a clonal TUBB-tagged TP53^-/-^ hepatocyte line is shown, highlighting the presence of a frameshift indel. **(b)** In a background of *TP53* mutations, frequent non-canonical mitotic spindles are observed in TUBB-tagged organoids. Arrows indicate mitotic spindles. **(c-d)** Representative snapshots of a time-lapse imaging experiment (5 min interval) in TUBB-tagged TP53^-/-^ organoids demonstrates the division of a polyploid hepatocyte with 3 nuclei through the formation of a multipolar spindle **(c)**, and the presence of a spindle with disorganized microtubules **(d). (e)** Quantification of the percentage of non-canonical mitotic spindles in both TUBB-tagged WT and TP53^-/-^ organoids. **(f)** Analysis of the type (in different colours) of non-canonical mitotic spindles. Each dot represents the percentage of the indicated type of aberration in one organoid, normalized over the total number of spindles quantified in that organoid. See **Supplementary Fig. 6** for representative images of each type of aberration. Data in **e** and **f** are the mean ± s.d. of n≥4 independent experiments and 3 clonal TUBB-tagged TP53^-/-^ lines from each of the 2 different donors analysed.

## Discussion

Here we show that NHEJ can consistently mediate precise gene insertion with high efficiency in human organoids derived from different tissues. The strategy outperforms HDR, even when combined with transient TP53 inhibition (**Fig. 1**). Importantly, since the knock-in efficiencies were assessed in wild-type primary human cells, they likely reflect the activities of these DNA repair pathways *in vivo.* These results are therefore more representative than the ones obtained in immortalized cell lines with highly divergent genomes.

By exploiting this mechanism of cellular DNA repair, we could target different loci in organoids derived from multiple healthy human tissues (**Supplementary Fig. 8**). This was not restricted to tagging highly expressed genes: we could also tag non-constitutively expressed markers for rare cell types (**Fig. 4d-e**). To facilitate this even further, we designed and tested different targeting plasmids that have a built-in selection strategy under the control of an independent promoter (**Supplementary Fig. 1**). This could aid selection of knock-in events at non-constitutively expressed loci, whose expression cannot be induced by transient differentiation as done in this study for *CHGA* and *MUC2*. In principle such a build-in selection strategy could be important in case of tagging genes with non-fluorescent tags (*e.g.* HA- or FLAG-tags).

This approach would be applicable to other human cell models, such as iPSC and ES-derived organoids. Although in these systems successful knock-in via conventional HDR approaches has already repeatedly been achieved^47-49^, use of an homology-independent-based strategy may increase the efficiency of knock-in generation.

Importantly, we could also easily create double knock-in lines (**Fig. 5e**). There has been previous attention to analysing the mode of division of hepatocytes to understand how the peculiar hepatocyte polarity in the liver is created and maintained^38^. These studies were so far confined to immortalized human liver cell lines, *e.g.* HepG2 cells, cultured in a 2D configuration^39,40^. This does not reflect the tissue architecture of the liver. Moreover, due to lack of robust knock-in approaches, these analyses were performed on fixed samples, which do not allow for real-time observation of the division process. Our double TUBB::mNEON;CDH1::tdTomato hepatocyte organoid reporter line overcomes these previous hurdles. For instance, we could dynamically observe the rare cases of formation and division of polyploid hepatocytes, which involves the transient formation of a membrane between the daughter cells that precedes a subsequent fusion event (**Fig. 6g**). Previous analyses of division of 2D-cultured primary murine hepatocytes suggested that polyploid hepatocytes can divide and reduce their ploidy by formation of multipolar spindles, so-called “reductive mitosis”^50,51^. Our observations confirm binucleated hepatocyte division as a rare event, but rather suggest maintenance of the polyploid state (**Fig. 6g**). These discrepancies could be due to differences in 2D *vs* 3D culturing of the cells, which has been shown to induce missegregation^41^ or alternatively due to mouse-human species differences.

We also show that CRISPR-HOT can be combined with gene knock-out. TP53 has been shown to be involved in regulating hepatocyte ploidy in homeostasis and upon damage in murine liver^52^. By mutating *TP53* in our TUBB-tagged hepatocyte lines, we found that loss of TP53 facilitates the division of polyploid cells and promotes abnormal mitotic spindle behaviour, which could result in aneuploidy. While aneuploidy is frequently associated with cancer, aneuploidy in the liver has been speculated to promote adaptation to liver injury^53^. However, the causal relation between *TP53* mutation and mitotic spindle aberrations suggests that the consequent aneuploidy may be linked to hepatocarcinogenesis.

In conclusion, CRISPR-HOT is a versatile strategy to achieve fast and efficient gene knock-in in human organoids. It will constitute a useful asset in studies requiring the generation of reporter lines, protein tagging, labelling of cellular structures, and lineage tracing experiments.

## Supporting information

Manuscript

## Acknowledgments

We would like to thank Helmuth Gehart and Amanda Andersson Rolf for kindly providing plasmids, Yotam Bar-Ephraïm and Jochem Bernink for help with FAC-sorting, Gabriella Darmasaputra and Matilde Galli for advice on hepatocyte ploidy, and Stieneke van den Brink for support with preparation of media components. The authors wish to thank the Hornung lab for providing the frame selectors and the targeting backbone. We sincerely acknowledge all the anonymous tissue donors. This work is part of the Oncode Institute which is partly financed by the Dutch Cancer Society and was funded by the gravitation program CancerGenomiCs.nl from the Netherlands Organisation for Scientific Research (NWO), MKMD grant (114021012) from Netherlands Organization for Scientific Research (NWO-ZonMw), and from Stichting Vrienden van Hubrecht 2012.10. BA was supported by a FEBS Long-term fellowship and is the recipient of a VENI grant (NWO-ALW 863.15.015).

## Author Contributions

Conceptualization: BA, DH; Methodology: BA, DH, HC; Software: BA, DH, RK, XZ; Formal Analysis: BA, DH, RK, XZ; Investigation: BA, DH, JB, RK, IJ, XZ; Resources: SCSL, ST, JvZ, HC; Data Curation: BA, DH, RK; Writing – Original Draft: BA, DH, HC: Writing – Review & Editing: BA, DH, JB, RK, XZ, HC; Visualization: BA, DH, RK; Supervision: HC; Project Administration: BA, DH; Funding Acquisition: BA, ST, JvZ, HC.

## Methods

### Human organoid culturing

For generation of human hepatocyte organoids, hepatocytes were isolated from human fetal livers obtained from Leiden University Medical Center. For generation of human liver ductal organoids, liver biopsies were obtained from patients undergoing surgery at Rotterdam University Medical Center. Isolation, generation and culture of human hepatocytes and human liver ductal organoids were performed as described in^30^ and^29^, respectively. This study is in compliance with all relevant ethical regulations regarding research involving human participants as decribed in^30^ and^29^. Human ileal tissues were obtained from the UMC Utrecht with informed consent of each patient in accordance with the Declaration of Helsinki and the Dutch law. Biopsies were taken from the normal mucosa of resected intestinal segments from a cancer patient. Human small intestinal crypts were isolated, processed and cultured as described previously^34^. Intestinal organoids were cultured in standard culture conditions (ENR)^30^. Differentiation was achieved by supplementing ENR medium with DAPT (10 μM, Sigma Aldrich) (for goblet cell enrichment) or by adding DAPT (10 μM, Sigma Aldrich) and the MEK inhibitor PD0325901 (500 nM; Sigma Aldrich) and withdrawing the p38 MAPK inhibitor SB202190, the TGF-β inhibitor A83-01, nicotinamide and Wnt-conditioned medium from the ENR medium (for enteroendocrine cells), as described in^8^.

### DNA plasmids

For the electroporation of human intestinal, hepatocyte and liver ductal organoids, different plasmids were used. The sgRNAs for the different targeted genes were cloned in the vector pSPgRNA (Addgene plasmid #47108), using the protocol described in^54^. Oligo sequences for each sgRNA are as follows:

AFP fwd 5’-CACCGTGCAGATTTCTCAGGCCTGT-3’

AFP rev 5’-AAACACAGGCCTGAGAAATCTGCAC-3’

CDH1 fwd 5’-CACCGAGGCGGCGAGGACGACTAG-3’

CDH1 rev 5’-AAACCTAGTCGTCCTCGCCGCCTC-3’

CHGA fwd 5’-CACCGCAGCTGCAGGCACTACGGCG-3’

CHGA rev 5’-AAACCGCCGTAGTGCCTGCAGCTGC-3’

KRT18 fwd 5’-CACCGCCAATGACACCAAAGTTCTG-3’

KRT18 rev 5’-AAACCAGAACTTTGGTGTCATTGGC-3’

KRT19 fwd 5’-CACCGCCCAGAGCCTGCTGCCTCAG-3’

KRT19 rev 5’-AAACCTGAGGCAGCAGGCTCTGGGC-3’

MUC2 fwd 5’-CACCGCATCTGGGGAGCGGGTGAGC-3’

MUC2 rev 5’-AAACGCTCACCCGCTCCCCAGATGC-3’

TUBB fwd 5’-CACCGAGGCGGCGAGGACGACTA-3’

TUBB rev 5’-AAACTAGGCCTCCTCTTCGGCCTC-3’

The frame selector plasmids containing a sgRNA that linearizes the donor targeting construct, the Cas9 and mCherry for detection of electroporated cells were obtained from Addgene (plasmids #66939, #66940, #66941). All targeting constructs (for both HDR and NHEJ) used in this study are shown in **Supplementary Fig. 1** and either designed and cloned in this study or obtained from^27^. For the HDR targeting constructs, the tags were flanked by 0.5 kb long homology arms for the specific genomic regions. The NHEJ targeting constructs designed in this study will be deposited in Addgene. To study the effect of transient TP53 inhibition on knock-in efficiency a plasmid encoding for a dominant negative form of TP53 (DN TP53) (Addgene #41856) was co-transfected with the respective plasmids.

### Organoid transfection and generation of knock-in lines

Transfection of human liver ductal organoids was performed using an electroporation-based protocol described in^31^. Briefly, the organoids were co-electroporated with the appropriate sgRNA plasmid, the frame selector plasmid and the targeting construct. Two days after electroporation, the transfected organoids were picked based on mCherry expression (from the frame selector plasmid) and dissociated into single cells with accutase (Life technologies cat# 00-4555-56) and re-plated in a BME^R^ (AMSBIO cat# 3533-005-02) drop. Growing knocked-in organoids (as judged based on fluorescence expression) were picked and used to establish clonal knocked-in organoid lines. Wnt-surrogate^55^ (0.15 nM, U-Protein Expression) and Rho kinase inhibitor (10 μM, Calbiochem Y27632) was added to the media whenever cells were at a single-cell stage to help recovery. For genotyping, organoids were lysed using a lysis buffer (0.1 M Tris-HCl pH 8.5, 0.2 M NaCl, 0.2% SDS, 0.05 M EDTA, 0.4 mg/ml ProteinaseK) and DNA was isolated by ethanol precipitation. PCR amplification followed by sequencing was used to confirm proper knock-in at the targeted locus. For analysis of knock-in efficiency, cells from the different samples were isolated from the BME^R^, digested into single cells with accutase and used for FACS analysis (FACS Aria, BD). DAPI was used to identify live cells. FlowJo software was used for data analyses.

Transfection of human hepatocyte organoids was performed by cuvette electroporation. Organoids were collected from 2 wells of a 12-well-plate and dissociated into single cells/small clumps using accutase. After 2 washes with PBS, organoids were resuspended in 150 μl of Opti-MEM and supplemented with the plasmids (10 μg total amount dissolved in 20 μl of PBS) and transferred to a 2 mm cuvette. Electroporation was performed using the following parameters on the NEPA21 electroporator: Poring Pulse (Voltage= 175 V, Pulse Length= 7.5 msec, Pulse Interval= 50 msec, Number of Pulse=2), Transfer Pulse (Voltage= 20 V, Pulse Length= 50 msec, Pulse Interval= 50 msec, Number of Pulse=5. Complete media supplemented with Rho kinase inhibitor (Calbiochem Y27632, final concentration 10 μM) was added to the cuvette. Cells were allowed to recover for 30 minutes after which they were seeded in BME^R^. Growing knocked-in organoids (as judged based on fluorescence expression) were picked and used to establish clonal human hepatocyte knock-in lines. For generation of the TP53 mutant hepatocyte organoid lines, TUBB-tagged organoids were electroporated with a TP53sgRNA-Cas9-GFP plasmid^11^ and clonal organoids with inactive TP53 were selected based on the addition of 10 μM Nutlin-3a (Cayman Chem #548472-68-0) to the media. Clonal lines were established and each line was genotyped to confirm the presence of bi-allelic TP53 mutations. Three clonal TP53^-/-^ lines from two TUBB-tagged hepatocyte lines originating from 2 different donors were generated. For knock-in efficiency analysis, hepatocyte organoids were similarly electroporated and 3 days after electroporation cells were recovered from the BME^R^ and used for FACS analysis (FACS Aria, BD). DAPI was used to identify live cells. FlowJo software was used for data analyses. Alternatively, to determine the preciseness of gene knock-in at two different loci, single transfected cells were FAC-sorted and seeded in BME^R^ to grow out single clonal organoids. Fluorescent images of entire 24-wells were acquired and number of fluorescent positive and negative organoids were quantified. Representative positive and negative organoids were picked and screened for their genotype by PCR amplification and subsequent sequencing.

Transfection of human intestinal organoids was performed by cuvette electroporation of a single well of a 12-well plate according to^36^ using a NEPA21 electroporator (Nepa Gene). Three to five days after electroporation, single transfected cells were FAC-sorted (FACS Aria, BD) based on transient expression of mCherry from the frame selector plasmid and seeded in BME^R^. For the generation of knock-in lines of constitutively expressed genes, single growing organoids were picked based on fluorescence to establish clonal lines. For the non-constitutively expressed genes, *i.e. MUC2* and *CHGA*, a differentiation pulse of 24 h with the respective differentiation media (compositions described above) was used. Single organoids showing a few fluorescent-positive induced cells were picked and transferred to expansion (ENR) medium for subsequent clonal line generation. In these clonal lines, differentiation to enrich for the specific differentiated intestinal cell types was performed by a switch from ENR to the respective differentiation media for 5 days.

All knock-in organoid lines from the different organs were confirmed for their genotype by PCR amplification and subsequent sequencing. Generation of knock-in lines at different loci was repeated multiple times (>3 experiments) from at least 2 independent donors for human liver ductal and hepatocyte organoids and 1 donor for human small intestinal organoids.

### Whole-mount immunofluorescence

For whole mount immunofluorescence staining, organoids were isolated from the BME^R^ using Cell recovery solution (Corning), fixed for 30 mins to 1 hour in 4% PFA, permeabilized and blocked by incubation in 0.2% Triton-X-100 and 5% normal donkey serum (Jackson Laboratories) in PBS for 1h at RT. Alternatively, staining was performed directly in the culture plate without dissociation of the BME^R^ drop and following the same procedure. Primary antibodies were incubated at 4°C overnight. Antibodies used in this study were: anti-AFP rabbit polyclonal 1:250 (Thermo Fisher #PA5-16658), anti-chromogranin A rabbit polyclonal 1:200 (labnet.com #LN1401487), anti-cleaved caspase-3 (Asp175) rabbit polyclonal 1:400 (Cell Signaling Technology #9661L), anti-E-cadherin mouse monoclonal 1:200 (BD bioscience #610182), anti-Keratin19 mouse monoclonal 1:200 (Cell Signalling #4558S), anti-Keratin20 mouse monoclonal 1:200 (Neomarkers #MS-377-S1), anti-MRP2 mouse monoclonal 1:100 (Abcam #ab3373), anti-Serotonin goat polyclonal 1:200 (Abcam #ab66047), anti-ZO-1 rabbit polyclonal 1:200 (life technology #402200). After washing with PBS, Alexa Fluor conjugated secondary antibodies were added for 1h at RT. Phalloidin-Atto 647N 1:1000 (Sigma #65906) was used for Actin staining. DAPI was used to counterstain nuclei. Organoids were mounted and imaged on an Sp8 confocal microscope (Leica). Images were processed using Photoshop CS4 or ImageJ software. Imaris software was used for images 3D-reconstruction and visualization purposes.

### Time-lapse imaging

For time-lapse microscopy imaging, organoids were passaged 2 days before and seeded on glass-bottom plates (Greiner #662892). Imaging was performed on a Sp8 confocal microscope (Leica) which was continuously held at 37°C and equipped with a culture chamber for overflow of 5% CO_2_. Imaging of the CDH1-, TUBB-, and double CDH1/TUBB-tagged hepatocyte organoids was performed for up to 72 hours, with acquisition intervals of 5 or 15 minutes and z-stack steps of 4 μm. For fluorescence detection, we used minimal amounts of excitation light. ImageJ software was used to assemble the videos and for image analysis. At least 2 different clonal lines were imaged and 4 independent imaging experiments were performed. Quantification of mitosis and types of non-canonical mitotic spindles was performed on the imaging of 15 organoids from the 3 different clonal lines derived from 2 different donors in independent experiments and was assessed by 2 researchers.

### Imaging analysis

For mitotic spindle analysis every spindle visible at any *xy*-plane was analysed. The coordinates of both ends of the spindle (spindle poles) were manually tracked over time. The spindle was tracked from the moment both ends were clearly separated (metaphase) to the moment just before the spindle disappeared (telophase). The 2D orientation of the line segment in between both ends was calculated for each time point, ignoring the *z* coordinate of the ends. For all spindles that were clearly rotating next to an internal lumen, the position on the edge of the lumen closest to the spindle was manually recorded. The rotation of mitotic spindles in the TUBB-tagged organoids was analysed using custom-written Python scripts (available upon request). Analysis of modes of cell division in the double CDH1/TUBB-tagged line was performed by tracing single mitotic events in ImageJ.

The cell movement in the CHD1-tagged organoids was traced manually. Based on labelling of the cell membrane, the position of the cell centroid was determined and was registered at every time point. The cells were selected such that every part of the organoid had its motion sampled. Arrows were then drawn from the position of a cell centroid at the first time point, pointing towards its position at the last time point. For presentation purposes, the arrows indicate their 2D orientation, ignoring the *z* position of the cells. Absolute average cell rotation degrees were calculated by averaging the movement of single cells within one organoid.

### Data visualization and statistical analysis

All data analyses, quantification, statistical evaluation was done in Rstudio. Data visualization was done in Rstudio using the ggplot2 package. Student’s *t*-tests were performed to compare different experimental groups and *p*-values are indicated. A *p* value of ≤ 0.05 was considered as the cut off for significance; * *p* ≤ 0.05; ** *p* ≤ 0.01; *** *p* ≤ 0.001.

## References

1. Clevers, H. Modeling Development and Disease with Organoids. Cell 165, 1586–1597 (2016).

2. Rossi, G., Manfrin, A. & Lutolf, M.P. Progress and potential in organoid research. Nat Rev Genet 19, 671–687 (2018).

3. Artegiani, B. & Clevers, H. Use and application of 3D-organoid technology. Hum Mol Genet (2018).

4. Farin, H.F. et al. Visualization of a short-range Wnt gradient in the intestinal stem-cell niche. Nature 530, 340–343 (2016).

5. Tetteh, P.W. et al. Replacement of Lost Lgr5-Positive Stem Cells through Plasticity of Their Enterocyte-Lineage Daughters. Cell Stem Cell 18, 203–213 (2016).

6. Barriga, F.M. et al. Mex3a Marks a Slowly Dividing Subpopulation of Lgr5+ Intestinal Stem Cells. Cell Stem Cell 20, 801–816 e807 (2017).

7. Kon, S. et al. Cell competition with normal epithelial cells promotes apical extrusion of transformed cells through metabolic changes. Nat Cell Biol 19, 530–541 (2017).

8. Beumer, J. et al. Enteroendocrine cells switch hormone expression along the crypt-to-villus BMP signalling gradient. Nat Cell Biol 20, 909–916 (2018).

9. Serra, D. et al. Self-organization and symmetry breaking in intestinal organoid development. Nature 569, 66–72 (2019).

10. Gehart, H. et al. Identification of Enteroendocrine Regulators by Real-Time Single-Cell Differentiation Mapping. Cell 176, 1158–1173 e1116 (2019).

11. Drost, J. et al. Sequential cancer mutations in cultured human intestinal stem cells. Nature 521, 43–47 (2015).

12. Drost, J. et al. Use of CRISPR-modified human stem cell organoids to study the origin of mutational signatures in cancer. Science 358, 234–238 (2017).

13. Schwank, G. et al. Functional repair of CFTR by CRISPR/Cas9 in intestinal stem cell organoids of cystic fibrosis patients. Cell Stem Cell 13, 653–658 (2013).

14. Shimokawa, M. et al. Visualization and targeting of LGR5(+) human colon cancer stem cells. Nature 545, 187–192 (2017).

15. Cortina, C. et al. A genome editing approach to study cancer stem cells in human tumors. EMBO Mol Med 9, 869–879 (2017).

16. Sugimoto, S. et al. Reconstruction of the Human Colon Epithelium In Vivo. Cell Stem Cell 22, 171–176 e175 (2018).

17. Essers, J. et al. Analysis of mouse Rad54 expression and its implications for homologous recombination. DNA Repair (Amst) 1, 779–793 (2002).

18. Hustedt, N. & Durocher, D. The control of DNA repair by the cell cycle. Nat Cell Biol 19, 1–9 (2016).

19. Haapaniemi, E., Botla, S., Persson, J., Schmierer, B. & Taipale, J. CRISPR-Cas9 genome editing induces a p53-mediated DNA damage response. Nat Med 24, 927–930 (2018).

20. Ihry, R.J. et al. p53 inhibits CRISPR-Cas9 engineering in human pluripotent stem cells. Nat Med 24, 939–946 (2018).

21. Schiroli, G. et al. Precise Gene Editing Preserves Hematopoietic Stem Cell Function following Transient p53-Mediated DNA Damage Response. Cell Stem Cell 24, 551–565 e558 (2019).

22. Betermier, M., Bertrand, P. & Lopez, B.S. Is non-homologous end-joining really an inherently error-prone process? PLoS Genet 10, e1004086 (2014).

23. Guo, T. et al. Harnessing accurate non-homologous end joining for efficient precise deletion in CRISPR/Cas9-mediated genome editing. Genome Biol 19, 170 (2018).

24. Auer, T.O., Duroure, K., De Cian, A., Concordet, J.P. & Del Bene, F. Highly efficient CRISPR/Cas9-mediated knock-in in zebrafish by homology-independent DNA repair. Genome Res 24, 142–153 (2014).

25. Suzuki, K. et al. In vivo genome editing via CRISPR/Cas9 mediated homology-independent targeted integration. Nature 540, 144–149 (2016).

26. Lackner, D.H. et al. A generic strategy for CRISPR-Cas9-mediated gene tagging. Nat Commun 6, 10237 (2015).

27. Schmid-Burgk, J.L., Honing, K., Ebert, T.S. & Hornung, V. CRISPaint allows modular base-specific gene tagging using a ligase-4-dependent mechanism. Nat Commun 7, 12338 (2016).

28. He, X. et al. Knock-in of large reporter genes in human cells via CRISPR/Cas9-induced homology-dependent and independent DNA repair. Nucleic Acids Res 44, e85 (2016).

29. Huch, M. et al. Long-term culture of genome-stable bipotent stem cells from adult human liver. Cell 160, 299–312 (2015).

30. Hu, H. et al. Long-Term Expansion of Functional Mouse and Human Hepatocytes as 3D Organoids. Cell 175, 1591–1606 e1519 (2018).

31. Artegiani, B. et al. Probing the Tumor Suppressor Function of BAP1 in CRISPR-Engineered Human Liver Organoids. Cell Stem Cell 24, 927–943 e926 (2019).

32. Okita, K. et al. An efficient nonviral method to generate integration-free human-induced pluripotent stem cells from cord blood and peripheral blood cells. Stem Cells 31, 458–466 (2013).

33. Broutier, L. et al. Culture and establishment of self-renewing human and mouse adult liver and pancreas 3D organoids and their genetic manipulation. Nat Protoc 11, 1724–1743 (2016).

34. Sato, T. et al. Long-term expansion of epithelial organoids from human colon, adenoma, adenocarcinoma, and Barrett’s epithelium. Gastroenterology 141, 1762–1772 (2011).

35. Fujii, M. et al. Human Intestinal Organoids Maintain Self-Renewal Capacity and Cellular Diversity in Niche-Inspired Culture Condition. Cell Stem Cell 23, 787–793 e786 (2018).

36. Fujii, M., Matano, M., Nanki, K. & Sato, T. Efficient genetic engineering of human intestinal organoids using electroporation. Nat Protoc 10, 1474–1485 (2015).

37. Yin, X. et al. Niche-independent high-purity cultures of Lgr5+ intestinal stem cells and their progeny. Nat Methods 11, 106–112 (2014).

38. Treyer, A. & Musch, A. Hepatocyte polarity. Compr Physiol 3, 243–287 (2013).

39. Wang, T., Yanger, K., Stanger, B.Z., Cassio, D. & Bi, E. Cytokinesis defines a spatial landmark for hepatocyte polarization and apical lumen formation. J Cell Sci 127, 2483–2492 (2014).

40. Lazaro-Dieguez, F. & Musch, A. Cell-cell adhesion accounts for the different orientation of columnar and hepatocytic cell divisions. J Cell Biol 216, 3847–3859 (2017).

41. Knouse, K.A., Lopez, K.E., Bachofner, M. & Amon, A. Chromosome Segregation Fidelity in Epithelia Requires Tissue Architecture. Cell 175, 200–211 e213 (2018).

42. Wang, M.J., Chen, F., Lau, J.T.Y. & Hu, Y.P. Hepatocyte polyploidization and its association with pathophysiological processes. Cell Death Dis 8, e2805 (2017).

43. Guidotti, J.E. et al. Liver cell polyploidization: a pivotal role for binuclear hepatocytes. J Biol Chem 278, 19095–19101 (2003).

44. Margall-Ducos, G., Celton-Morizur, S., Couton, D., Bregerie, O. & Desdouets, C. Liver tetraploidization is controlled by a new process of incomplete cytokinesis. J Cell Sci 120, 3633–3639 (2007).

45. Aylon, Y. & Oren, M. p53: guardian of ploidy. Mol Oncol 5, 315–323 (2011).

46. Vogel, C., Kienitz, A., Hofmann, I., Muller, R. & Bastians, H. Crosstalk of the mitotic spindle assembly checkpoint with p53 to prevent polyploidy. Oncogene 23, 6845–6853 (2004).

47. Zhang, J.P. et al. Efficient precise knockin with a double cut HDR donor after CRISPR/Cas9-mediated double-stranded DNA cleavage. Genome Biol 18, 35 (2017).

48. Mitzelfelt, K.A. et al. Efficient Precision Genome Editing in iPSCs via Genetic Co-targeting with Selection. Stem Cell Reports 8, 491–499 (2017).

49. Zhang, J.Z. et al. A Human iPSC Double-Reporter System Enables Purification of Cardiac Lineage Subpopulations with Distinct Function and Drug Response Profiles. Cell Stem Cell 24, 802–811 e805 (2019).

50. Duncan, A.W. et al. The ploidy conveyor of mature hepatocytes as a source of genetic variation. Nature 467, 707–710 (2010).

51. Duncan, A.W. et al. Frequent aneuploidy among normal human hepatocytes. Gastroenterology 142, 25–28 (2012).

52. Kurinna, S. et al. p53 regulates a mitotic transcription program and determines ploidy in normal mouse liver. Hepatology 57, 2004–2013 (2013).

53. Duncan, A.W. et al. Aneuploidy as a mechanism for stress-induced liver adaptation. J Clin Invest 122, 3307–3315 (2012).

54. Ran, F.A. et al. Genome engineering using the CRISPR-Cas9 system. Nat Protoc 8, 2281–2308 (2013).

55. Janda, C.Y. et al. Surrogate Wnt agonists that phenocopy canonical Wnt and beta-catenin signalling. Nature 545, 234–237 (2017).

