## Supplementary material for "Fast and efficient generation of knock-in human organoids using homology-independent CRISPR/Cas9 precision genome editing": Manuscript

Supplementary information includes 3 supplementary videos (**Supplementary Video 1-3**) and 8 supplementary figures (**Supplementary Figures 1-8**).

**Supplementary Video 1.** Representative examples of time-lapse imaging of TUBB-tagged WT human hepatocyte organoids from two donors.

**Supplementary Video 2.** Representative examples of time-lapse imaging of double TUBB::mNEON;CDH1::tdTomato WT human hepatocyte organoids.

**Supplementary Video 3.** Representative examples of time-lapse imaging of TUBB-tagged TP53$^{-/-}$ human hepatocyte organoids from two donors.

### Supplementary figure 1

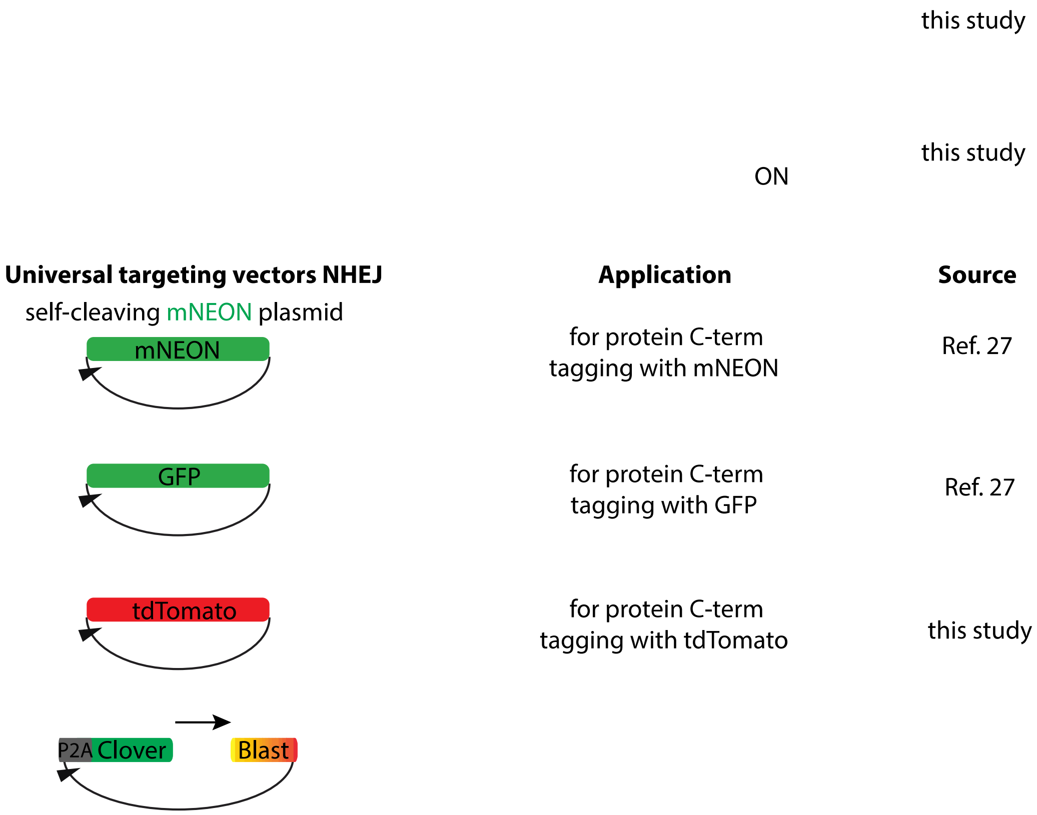


**Supplementary Figure 1: Schematic overview of NHEJ and HDR targeting plasmids used in this study.** List of plasmids used or generated in this study for gene tagging, including a schematic map of the plasmid, the application and the plasmid source.

### Supplementary figure 2

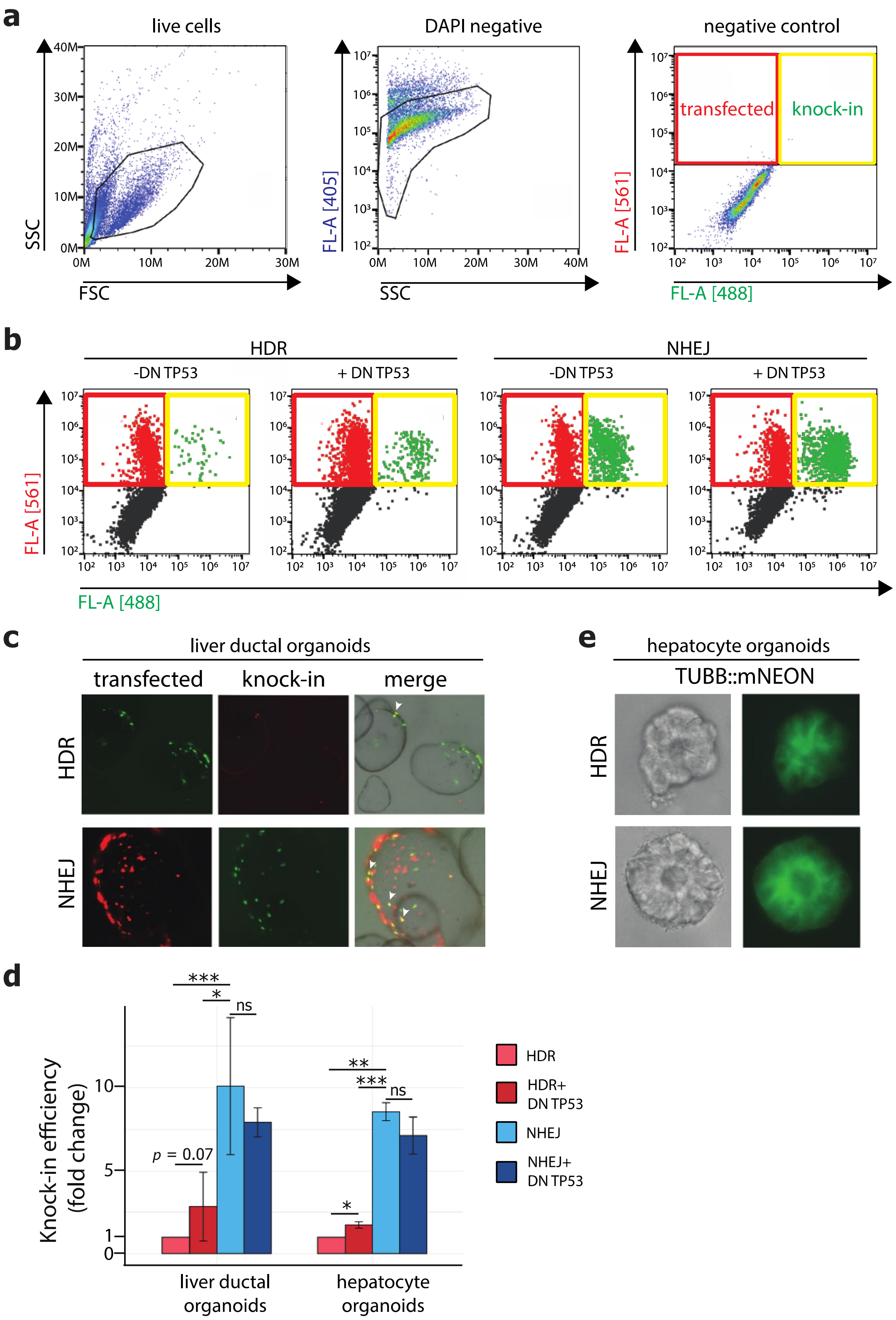

**Supplementary Figure 2: FACS analysis of knock-in efficiencies mediated by NHEJ or HDR.** **(a)** Gating strategy and representative plot of how transfected and knocked-in cells were defined based on the negative control (non-transfected cells). **(b)** Representative FACS plots of HDR- and NHEJ-mediated knock-in in presence or absence of a dominant negative form of TP53 (DN TP53) when targeting the *TUBB* locus in human hepatocyte organoids. **(c)** Representative images of electroporated human liver ductal organoids for both HDR- and NHEJ-mediated in-frame knock-in of the respective tag at the *KRT19* locus, showing knocked-in cells overlapping within the transfected cells. **(c)** Representative plot for HDR-mediated in-frame knock-in of tdTomato at the KRT19 locus in transfected cells based on transient GFP expression. **(d)** Bar plot showing fold changes of knock-in efficiency between HDR and NHEJ in presence or absence of a dominant negative form of TP53 (DN TP53) in human liver ductal organoids (targeting the *KRT19* locus) and human hepatocyte organoids (targeting the *TUBB* locus). **(e)** Representative brightfield and fluorescent images of HDR- and NHEJ-mediated knocked-in mNEON at the *TUBB* locus in human hepatocyte organoids.

### Supplementary figure 3

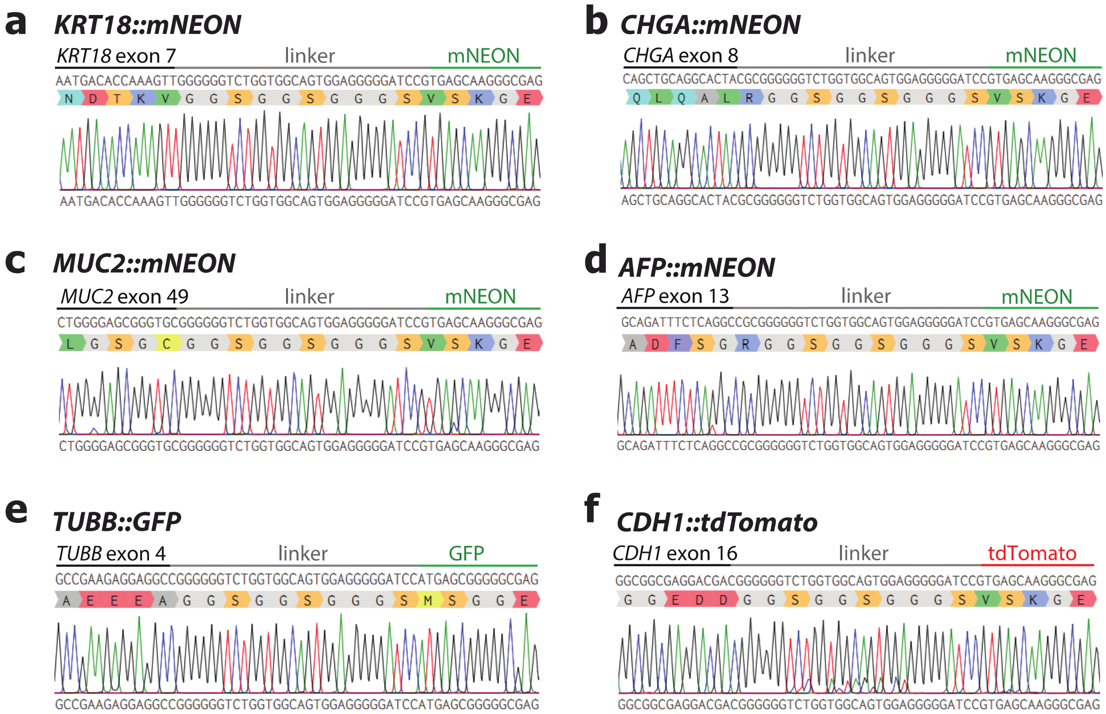


**Supplementary Figure 3: Representative sequencing results of generated human knock-in organoid lines in this study.** Sequencing results from clonal organoid lines spanning the insertion site for KRT18::mNEON **(a)**, CHGA::mNEON **(b)**, MUC2::mNEON **(c)**, AFP::mNEON **(d)**, TUBB::GFP **(e)**, CDH1::tdTomato **(f)**. For every targeted locus, sequencing results from 1 of the tagged lines is shown.

#### Supplementary figure 4

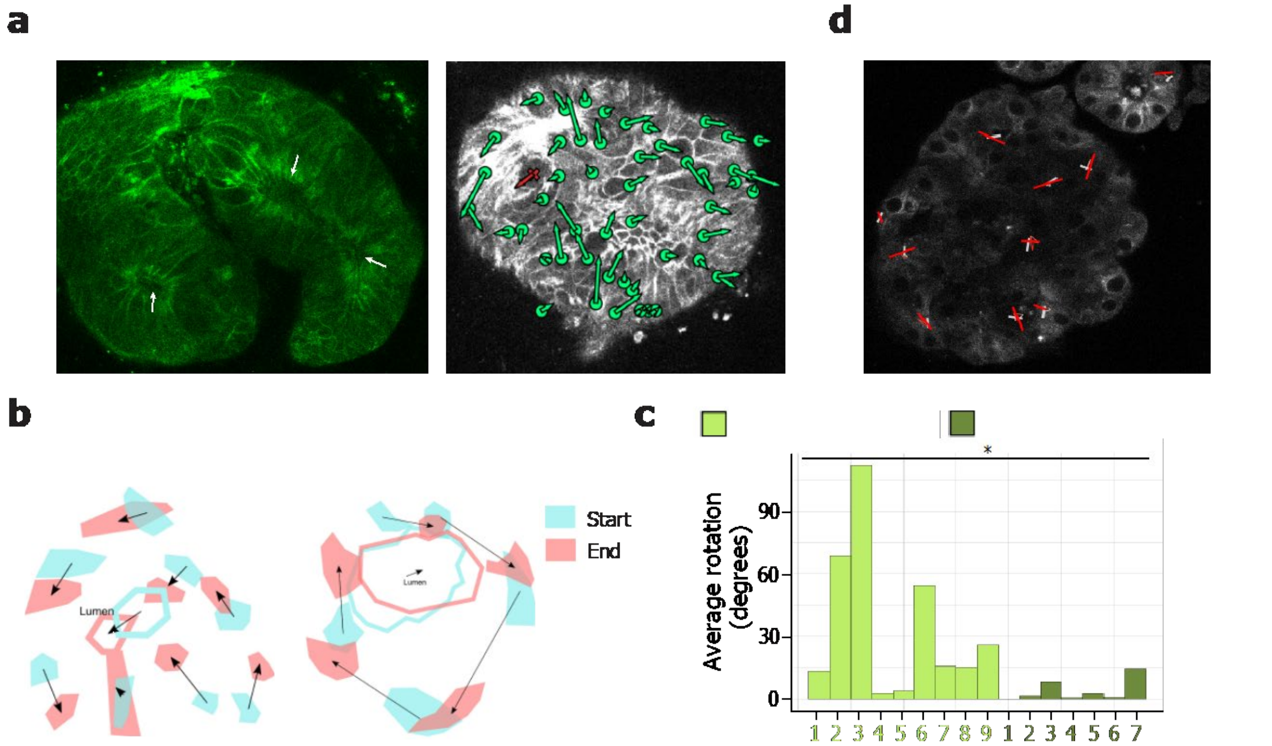


**Supplementary Figure 4: CDH1-tagged and TUBB-tagged human hepatocyte organoids as a tool for tracing cell movement and mitotic spindle dynamics. (a)** (left) Tagging of CDH1 reveals that hepatocytes tend to form rosette structures around lumina (indicated by the white arrows). Analysis of cell movement in CDH1::mNEON human hepatocyte organoids. (right) Representative example of an CDH1::mNEON human hepatocyte organoid in which individual cell movements were traced based on changes of cell centroid positioning over time. The green dots represent the initial position of the cell centroid and length and orientation of each green arrow represent individual cell centroid movement from begin to end of the experiment. The red line indicates lumen movement. (t = 9 hours, 45 min intervals). **(b)** Representative examples of tracing of cell movement based on changes of cell centroid positioning over time in CDH1::mNEON (left) and CDH1::tdTomato (right) human hepatocyte organoids. Initial and final positioning and cell shape outline are represented in blue and pink, respectively, and cell movements are indicated by arrows. Similarly, the initial and final position of the lumen is outlined and its movement is visualized. Note that hepatocytes tend to rotate around the lumen. **(c)** Bar plot showing the average cell rotation for individual organoids with a lumen (light green) or without a lumen (dark green). Note that in organoids in which there is no lumen cells tend to rotate less. as previously mentioned, organoids with a lumen tend to rotate. **(d)** Example of the determination of mitotic spindle dynamics in TUBB::GFP human hepatocyte organoids. The mitotic spindle orientation was traced over time by marking the initial (gray) and final (red) spindle orientation. The thin line indicates the average position of the spindle poles at every time point between the two.

### Supplementary figure 5

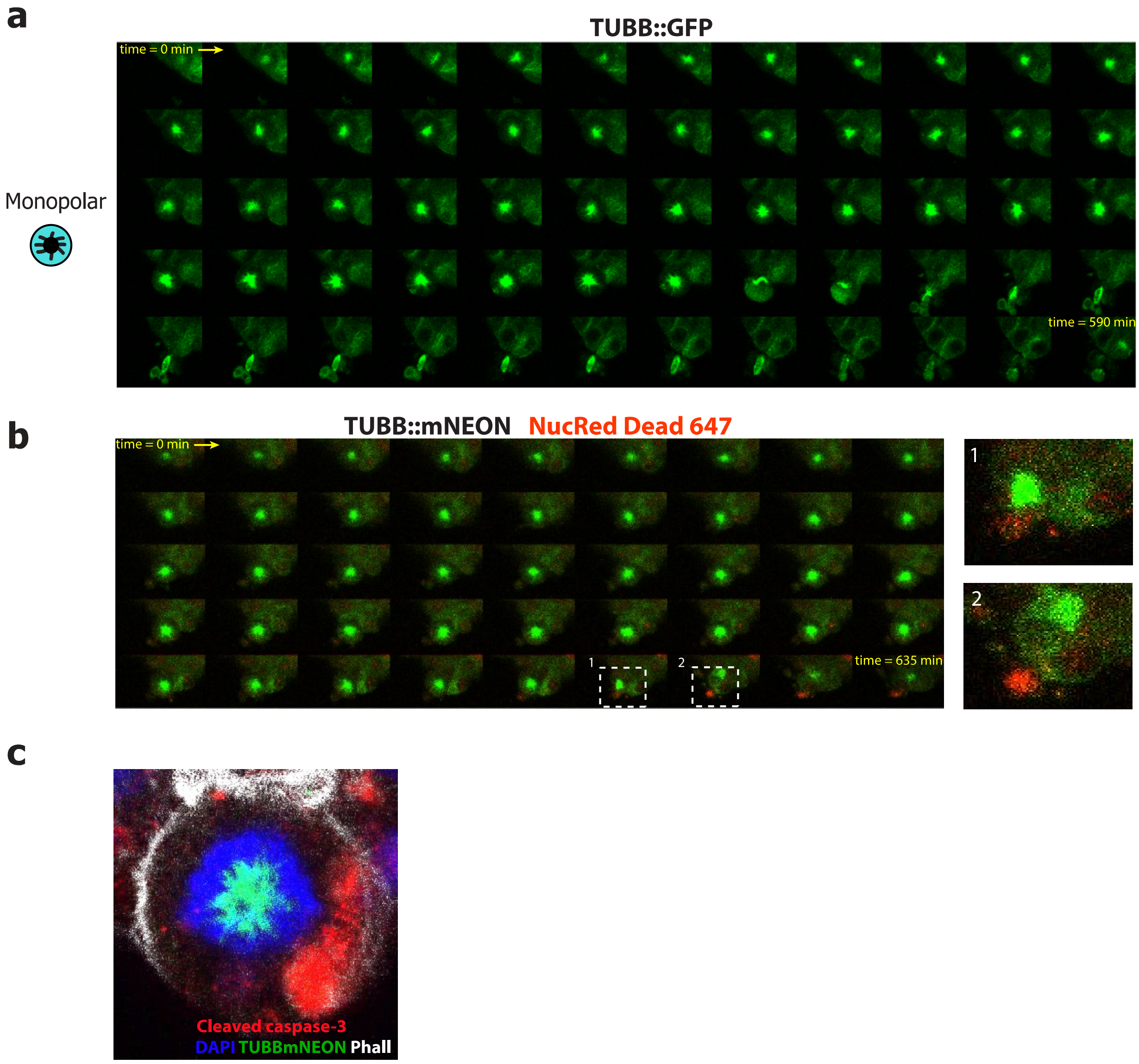


**Supplementary Figure 5: Characterization of cells with a monopolar spindle in TUBB-tagged human hepatocyte organoids. (a)** Representative snapshots of a time-lapse experiments showing the formation and fate of monopolar spindles in TUBB::GFP human hepatocyte organoids, resulting in cell bursting. **(b)** Representative snapshots of a time-lapse experiments in which organoids were incubated with NucRed Dead 647 confirming that cells with a monopolar spindle die. **(c)** Cells with a monopolar spindle stain positive for cleaved caspase-3.

#### Supplementary figure 6

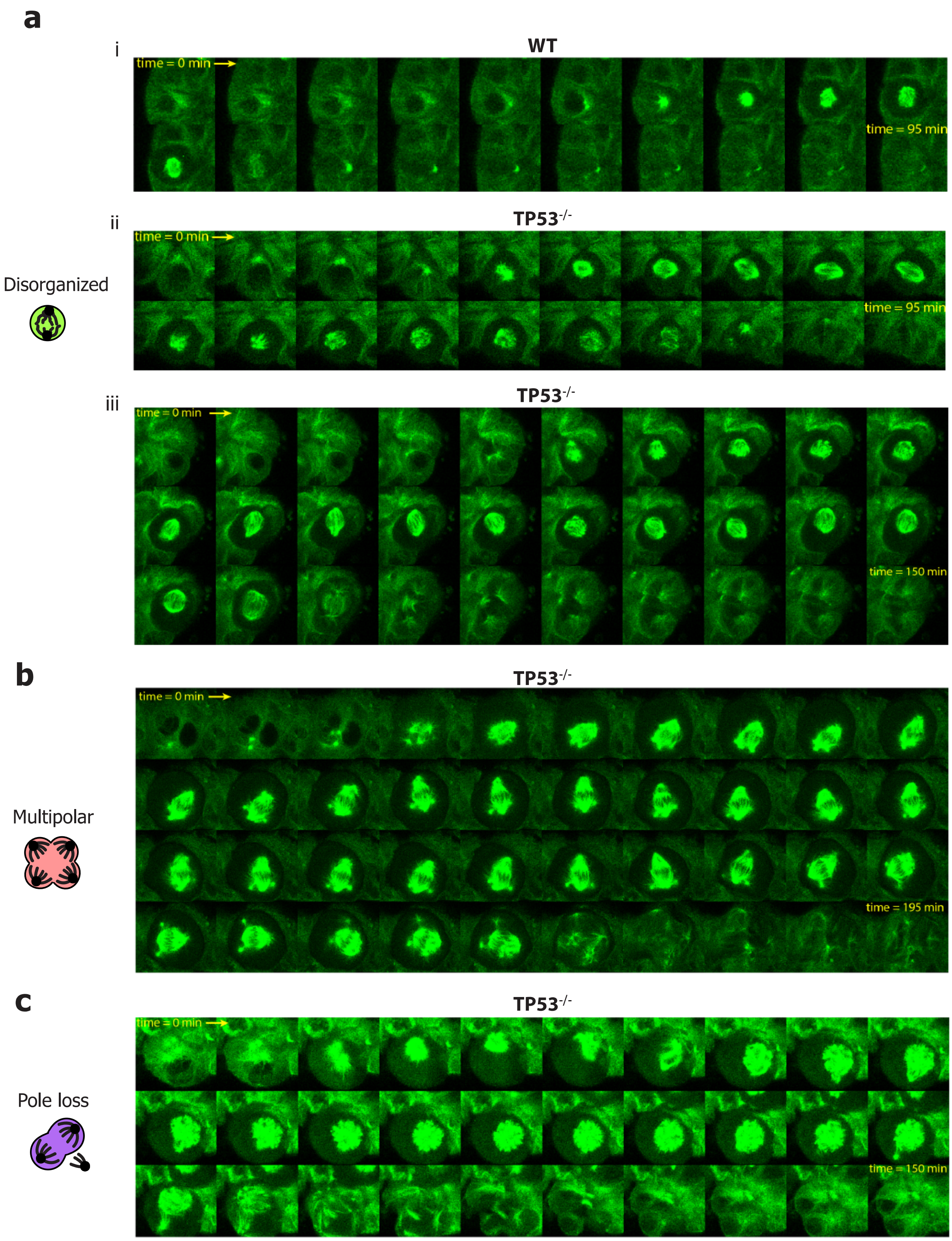


**Supplementary Figure 6: Types of non-canonical mitotic spindles in WT and TP53$^{-/-}$ human hepatocyte organoids. (a)** Examples of mitotic spindles with disorganized microtubules in a WT (i) and TP53$^{-/-}$ (ii, iii) background. **(b)** Loss of *TP53* causes frequent formation of multipolar spindles. **(c)** Example of a non-canonical mitotic event where a spindle pole is lost in a TP53$^{-/-}$ background.

### Supplementary figure 7

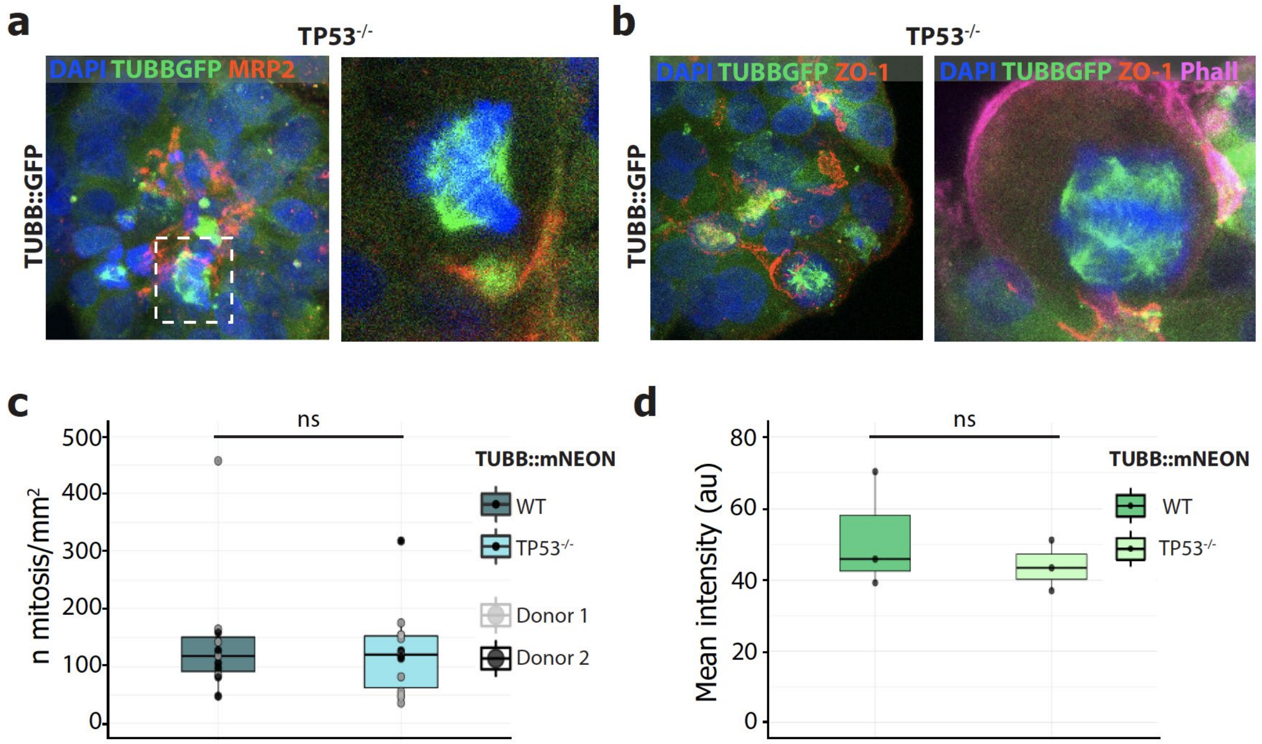


**Supplementary Figure 7: Characterization of the impact of TP53 mutation on the phenotype of human hepatocyte organoids.** Representative examples of whole-mount (left) and a focal plane (zoomed on the right) staining of TUBB::GFP TP53$^{-/-}$ organoids showing the intactness of the MRP2-marked bile canalicular network **(a)** and maintenance of hepatocyte polarity as marked by ZO-1, associated with a cell division with a multipolar spindle **(b)**. **(c)** Box-plot showing the amount of mitosis in WT and TP53$^{-/-}$ TUBB-tagged hepatocytes. Each dot represents mitosis quantification in one single organoid. Mitosis in organoids from 2 donors were quantified. **(d)** Box-plot showing similar TUBB::mNEON fluorescence intensity between WT and TP53$^{-/-}$ TUBB::mNEON hepatocyte organoids.

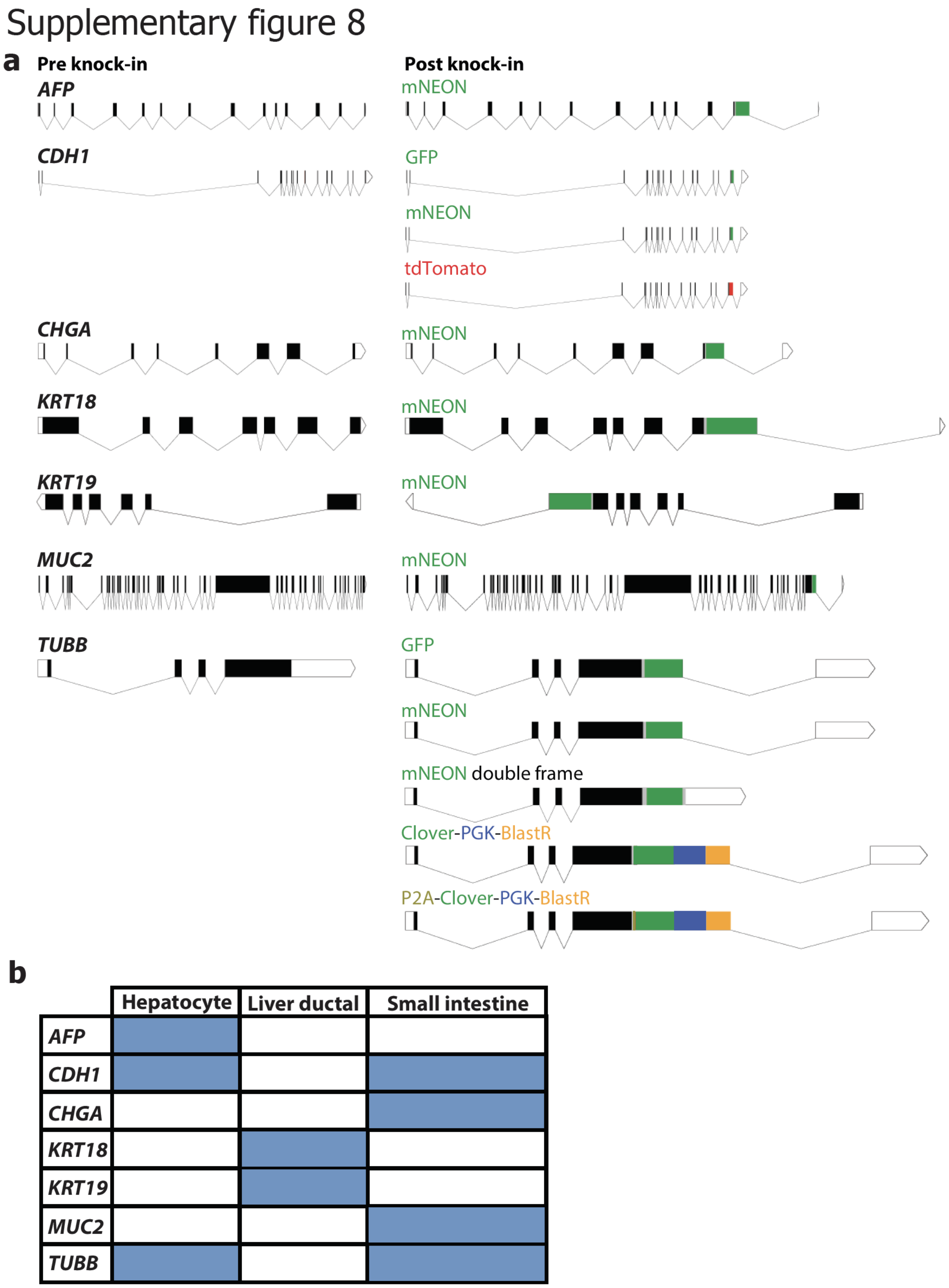


**Supplementary Figure 8: Schematic overview of all targeted loci in this study. (a)** Schematic representation of all the genes that have been tagged in this study showing a schematic of the gene pre and post knock-in with all the different tags. Black boxes represent exons, lines represent introns. **(b)** Overview of all the genes that have been tagged in the different organoid systems in this study.
